# Protein Interaction Screen on Peptide Matrix (PRISMA) reveals interaction footprints and the PTM-dependent interactome of intrinsically disordered C/EBPβ

**DOI:** 10.1101/238709

**Authors:** Gunnar Dittmar, Daniel Perez Hernandez, Elisabeth Kowenz-Leutz, Marieluise Kirchner, Günther Kahlert, Radoslaw Wesolowski, Katharina Baum, Maria Knoblich, Arnaud Muller, Jana Wolf, Ulf Reimer, Achim Leutz

## Abstract

CCAAT enhancer binding protein beta (C/EBPβ) is a pioneer transcription factor that specifies cell differentiation. C/EBPβ is intrinsically unstructured, a molecular feature common to many proteins involved in signal processing and epigenetics. The structure of C/EBPβ differs depending on alternative translation start site usage and multiple post-translational modifications (PTM). Mutation of distinct PTM sites in C/EBPβ alters designated protein interactions and cell differentiation, suggesting a C/EBPβ PTM indexing code determines epigenetic outcomes. Herein, we systematically explored the interactome of C/EBPβ using an array of spot-synthesised C/EBPβ-derived linear tiling peptides with and without PTM, combined with mass spectrometric proteomic analysis of protein interactions. We identified interaction footprints of ~1300 proteins in nuclear cell extracts, many with chromatin modifying, remodelling and RNA processing functions. The results suggest C/EBPβ acts as a multi-tasking molecular switchboard, integrating signal-dependent modifications and structural plasticity to orchestrate interactions with numerous protein complexes directing cell fate and function.

**Highlights:**

- Peptide array based interaction proteomics map SLiM and PTM dependent C/EBPβ interactome
- Novel links between C/EBPβ, RNA processing, transcription elongation, MLL, NuRD were revealed
- C/EBPβ structure organizes modular hub function for gene regulatory machinery
- PRISMA is suitable to resolve protein interactions and networks based on intrinsically disordered proteins

## Introduction

CCAAT enhancer binding proteins (C/EBPα, β, δ, ε) are basic leucine zipper transcription factors that regulate chromatin structure and gene expression by dimerisation and binding to cis-regulatory, palindromic 5’ATTGC·GCAAT3’, or quasi-palindromic DNA sites in gene enhancers and promoters. Prototypic C/EBPβ is widely expressed, highly regulated at the post-transcriptional level, and integrated in many signalling events communicating extracellular cues to epigenetic changes, examples of which include adipogenesis, haematopoiesis, innate immunity, female fertility, skin function, apoptosis, and cellular senescence (Nerlov, 2008; Rodier and Campisi, 2011; Tsukada et al., 2011).

In early haematopoiesis and adipogenesis, C/EBPβ acts as a pioneering factor that orchestrates complex steps in cell fate commitment (Kajimura et al., 2009; Lichtinger et al., 2012; Muller et al., 1995; Ness et al., 1993; Siersbaek et al., 2011). C/EBPβ communicates with numerous other transcription factors, co-factors, histone modifiers, and chromatin remodelling complexes to alter the susceptibility of chromatin to the gene regulatory machinery in lymphoid-myeloid trans-differentiation, and accelerates acquisition of the induced pluripotent state by the Yamanaka set of reprogramming transcription factors (Di Stefano et al., 2016; Kowenz-Leutz and Leutz, 1999; Stoilova et al., 2013; Xie et al., 2004). The chromatin and gene regulatory functionality of C/EBPβ is linked to distinct regions and post-translational modifications (PTM) that suspend auto-inhibition, direct the activity of C/EBPβ, and regulate recruitment of chromatin remodellers and writers of histone modifications that alter the structure of chromatin (Kowenz-Leutz and Leutz, 1999; Kowenz-Leutz et al., 2010; Kowenz-Leutz et al., 1994; Lee et al., 2010b; Mo et al., 2004; Pless et al., 2008; Siersbaek et al., 2011).

The complexity and diversity of C/EBPβ activities in various cell lineages raises the question of how a single transcription factor can participate in a multitude of regulatory events. C/EBPβ functions are controlled by extracellular signalling cascades involving receptor tyrosine kinases, cytokine receptors, Ras GTPases, MAP kinases, and cAMP and SMAD signalling, and by more complex conditions such as metabolic adaptation, inflammation, senescence or stress responses (Nerlov, 2008; Rodier and Campisi, 2011; Tsukada et al., 2011). Previous research suggested that the combinatorial outcome of post-transcriptional and post-translational modifications in conjunction with intrinsic structural plasticity enables the C/EBPβ protein to adopt a plethora of context- and signal-dependent states that facilitate a variety of interactions (Leutz et al., 2011; Nerlov, 2008; Tsukada et al., 2011).

Post-transcriptional modification of C/EBPβ generate three isoforms (LAP*, LAP, LIP) by alternative translation initiation of the single exon C/EBPβ transcript. Since consecutive C/EBPβ start sites are positioned in the same reading frame, the isoforms vary in their gene regulatory N-terminal extensions but retain the same C-terminal dimerisation and DNA-binding bZip domain (Wethmar et al., 2010). The diversity of C/EBPβ isoforms is further increased by numerous PTM of amino acid side chains; in addition to phosphorylation of serine, threonine, and tyrosine residues, lysine acetylation and methylation also occurs, as does methylation of arginine. Enzymes responsible for PTM of C/EBPβ include CARM1/PRMT4, G9A/EHMT2, and CREBBP/KAT3A, all of which serve as epigenetic histone code writers. The decoration of C/EBPβ by PTM alters its capacity to engage in protein-protein interactions (PPI) and to direct cell fate, suggesting that the signal-dependent C/EBPβ modification index reflects integration of various upstream signalling events to adjust its interactome and to determine its gene regulatory and epigenetic capacity (Leutz et al., 2011). The combination of translational and post-translational modifications may thus encrypt the dynamic interactome, with C/EBPβ as the keystone for a wide range of functional outcomes (Lee et al., 2010b; Sebastian et al., 2005; Sterneck et al., 1997; Stoilova et al., 2013).

The C-terminal third of C/EBPα, β, δ, ε contains highly conserved DNA binding and basic leucine zipper domains (bZip) that may dimerise within an extended trans-regulatory bZip network including C/EBP, AP-1, and ATF transcription factors. The N-terminal two-thirds of C/EBP primary sequences are predicted to be intrinsically disordered regions (IDRs). Phylogenetic analysis nevertheless suggests that the C/EBP N-terminus also contains several highly conserved short peptide regions (CRs) that are enriched in amino acids with hydrophobic and bulky side chains. These CRs are discontinuous and separated by less conserved and family-specific regions of low complexity (LCRs) characterised by a predominance of small and polar amino acids (Leutz et al., 2011; Tsukada et al., 2011). Experimental studies involving a large number of deletions and CR/IDR shuffling mutants suggested a highly modular, context-dependent functionality of N-terminal C/EBPβ CRs (Kowenz-Leutz et al., 1994; Lee et al., 2010b; Leutz et al., 2011; Williams et al., 1995). Screening for interaction partners using N-terminally-derived C/EBPβ peptides further supports the notion that many C/EBPβ interactions may occur in a modular and dynamic fashion that rely on molecular recognition features (MoRFs) contained in short linear peptide motifs (SLiMs), in combination with adjacent PTM (Dunker et al., 1998; Leutz et al., 2011; Tompa et al., 2014; van der Lee et al., 2014; Wright and Dyson, 1999).

Comprehension of the dynamics and context-dependent interactions between structurally flexible SLiMs and various partner proteins is an emerging hallmark of signal transmission and key to understanding the regulation of chromatin structure and gene regulation (Minde et al., 2017; Tompa et al., 2014; van der Lee et al., 2014; Wright and Dyson, 2015). Deciphering the functionality of individual protein regions and PTMs can be challenging when using full-length proteins due to functional redundancy and compensatory effects. Based on previous observation of the modular structure and functionality of C/EBPβ peptide regions, we developed a workflow to systematically explore the interactome by deconstructing C/EBPβ using a solid-phase array of small linear peptides. Briefly, synthetic unmodified and PTM-modified C/EBPβ tiling peptides were immobilised on a solid matrix that was subsequently used for affinity enrichment of soluble proteins and protein complexes from nuclear extracts. Proteins interacting with the C/EBPβ peptide matrix were then identified and quantified by mass spectrometry. The assay, termed **PR**otein Interaction **S**creen on peptide **Ma**trices (PRISMA), revealed hundreds of C/EBPβ- and PTM-specific protein interactions, and facilitated the identification of dozens of protein complexes as potential interaction partners that left footprints on C/EBPβ-derived peptides. Based on comparison with other affinity enrichment approaches, conventional immunoblotting analysis, and co-occurrence, several protein interactions and protein complexes were predicted and subsequently confirmed experimentally.

The linear C/EBPβ peptide interactome thus provides a repository of high molecular resolution data for protein interactions of an intrinsically disordered protein. The PRISMA approach serves as a basis to explore gene regulatory and PTM-modulated C/EBPβ functions, and can assist the rational design of mutant proteins for interaction studies. The workflow is applicable to other regulatory proteins and transcriptional regulators with similar structural features such as Myc and the various Hox proteins, and may help to discern highly complex and dynamic transcription factor functions that arise through IDR- and PTM-regulated interactions.

## Results & Discussion

### The C/EBPβ peptide matrix

The occurrence of novel PTMs on endogenous C/EBPβ was investigated by mass spectrometry of C/EBPβ immunoprecipitates derived from the human anaplastic lymphoma cell line SU-DHL1 (Anastasov et al., 2010; Jundt et al., 2005). Over 90 PTMs were identified (**Supplemental Table 1**) which, combined with published data, suggested that more than 130 PTMs may occur on this protein, as summarised in **Figure 1A**. To systematically explore the linear region and PTM specific C/EBPβ interactome, we designed a solid matrix consisting of immobilised peptides spanning the entire primary sequence of rat C/EBPβ (297 amino acids), as schematically depicted in **Figure 1B** and detailed in **Supplemental Table 2**. To cover all linear binding regions of the entire CEBPβ protein sequence, tiling peptides of 14 amino acids in length with an offset of mostly four amino acids were spot-synthesised on a cellulose acetate matrix using Fmoc synthesis. Since PTMs were previously shown to impact on CEBPβ protein interactions and functionality, PTM peptides with S/T/Y-phosphorylation, K-acetylation, K-, R-methylation, and R-citrullination were included in the screen matrix. In total, the solid matrix contained 203 immobilised peptides, covering known and potential post-translational side chain modifications (**Supplemental Table 2**).

**Figure 1:**
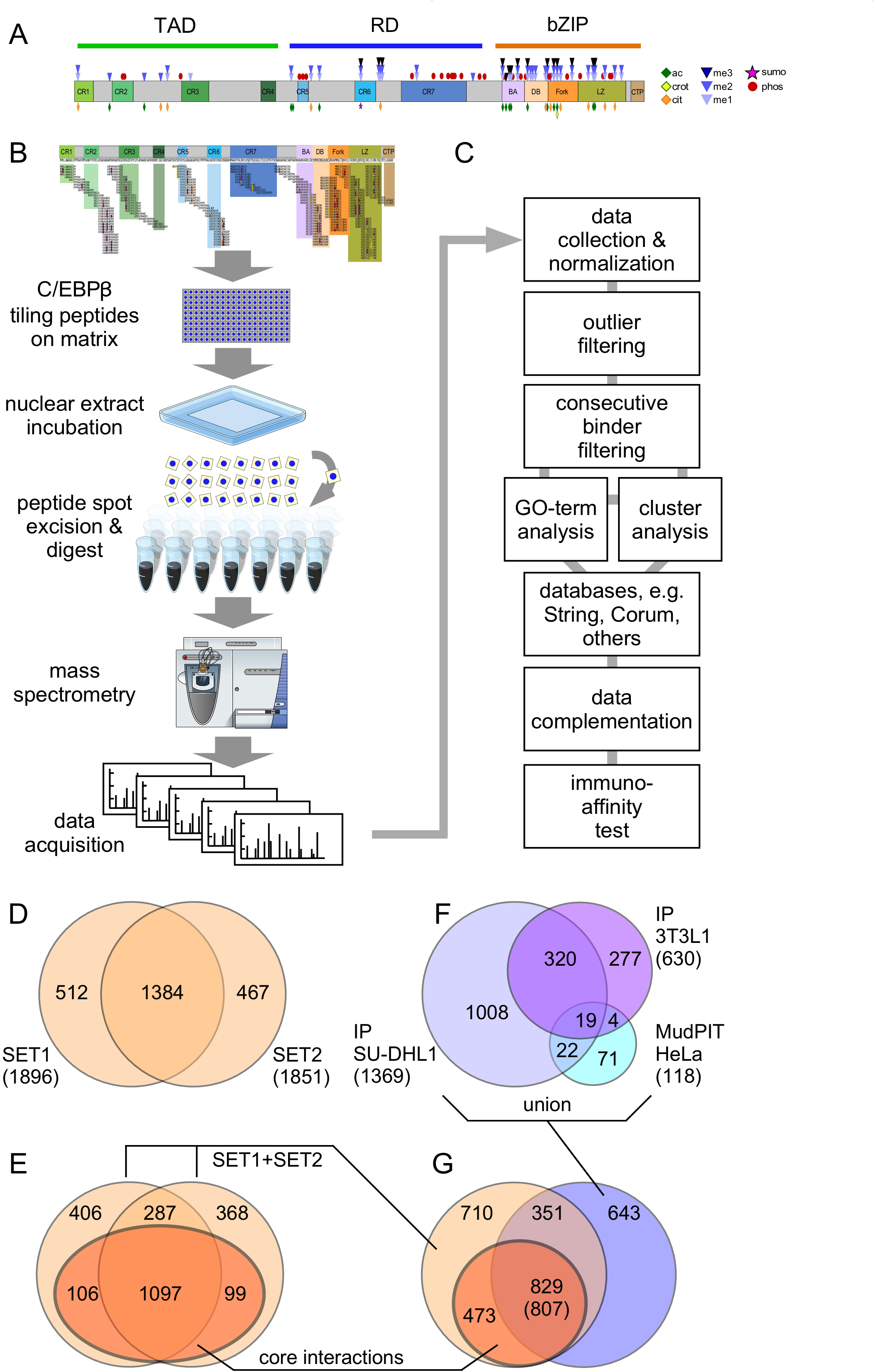
Outline of PRISMA screen and comparison of data. **A**. Schematic representation of C/EBPβ and the distribution of known post-translational modifications (PTMs). Conserved regions (CRs) are depicted in colour, while intrinsic disordered regions (IDR) are shown in grey. **B**. Schematic description of the workflow of protein binding and data acquisition. **C**. Workflow of data and proteomic analysis. **D**. Overlap between the two replicates (SET1, SET2) of the PRISMA screen. **E**. Number of proteins in the two PRISMA datasets that show consecutive peptide binding (core interactions, dark amber). **F**. Overlap of three affinity purification-based datasets of C/EBPβ interactors as described in the literature and combined with data obtained from a proteomic interaction screen in SU-DHL1 cells. Overlaps were determined using IP SU-DHL1 as reference dataset and thus the numbers add up to the size of this set only (see Material and Methods). **G**. Overlap of the PRISMA-derived C/EBPβ interactor datasets from E (union of SET1 and SET2, light amber) with core interactions (dark amber) from the union of the three datasets from F (blue). Overlaps are given using the PRISMA-derived data as reference datasets. The overlap count using the union of the datasets from F as a reference dataset is denoted in brackets.

### The C/EBPβ PRISMA screen

To examine the linear CEBPβ interactome, two replicates of the peptide matrix were incubated with HeLa nuclear extracts (**Figure 1B**). Individual peptide spots were excised and bound proteins proteolytically digested and analysed by high-resolution mass spectrometry. In total, 406 analytical mass spectrometric 1 h runs were performed (approximately 17 days of measurement) and spectra were interpreted automatically using the MaxQuant software package.

Enrichment efficiency by PRISMA was examined by comparing the PRISMA intensity distribution with the total proteome of the nuclear extract. Approximately 5100 proteins were identified and quantified in the nuclear cell extract using the intensity-based absolute quantification (iBaq) method (Schwanhausser et al., 2011). The copy number of proteins in the nuclear cell extract and the number of proteins bound in the PRISMA screen spans six orders of magnitude, suggesting that the identified binders are not biased towards highly abundant proteins (Smits et al., 2013) (**Supplemental Figure 1A**). Furthermore, the distribution of peptide/protein interactions was not attributed to physico-chemical parameters of the peptides, as evidenced by comparison of the accumulated interaction intensities with peptide hydrophobicity (gravy index) and isoelectric points (**Supplemental Figure 1B, C**), which were determined for all non-modified PRISMA peptides. These results suggest that the PRISMA peptides on the matrix retained their specific protein-binding properties.

### Data processing of C/EBPβ peptide binding proteins

An initial inspection of the two replicate datasets revealed signal intensity variation of the interacting proteins. The two datasets were therefore integrated to increase robustness, as outlined in **Figure 1C** and as explained in the Materials and Methods. The main source of variation arose from proteins that were identified only once on different peptide spots. This led to single (low confidence) and double (high confidence) identification categories for each peptide. The signal intensity for each protein was then normalised between 0 and 1 across all 203 matrix peptides. Individual peptides displayed large differences in binding partner profiles, further indicating the specificity of the interactions, because random binding would be expected to result in a more equal distribution of interacting proteins. Further analysis of the data showed that ~25% of the identified proteins bound to multiple C/EBPβ peptides across the array with low but varying intensity. While these proteins may promiscuously bind to many distant peptides, some sections of the array showed much higher signals. In order to minimise noise from background binding, signal intensities for each protein were filtered for binding above 90% of the protein’s signal distribution (outlier filtering), removing all signals below this threshold. Another filtering criterion for discriminating the most robust interactors was based on the consecutive binding behaviour of tiling peptides of the C/EBPβ primary sequence. The rational here is that by shifting the sequence of tiling peptides by four amino acids, some of the SLiMs and fractions thereof are included in more than one peptide. This may generate maximal binding signals for peptides containing optimal SLiM and adjacent supporting amino acids, and attenuate signals from neighbouring peptides in which the particular SLiM is shifted or only partially included. We used this predicted binding behaviour to stringently filter the dataset further to remove all proteins that failed the consecutive binding criterion (**Figure 1D, E**; see also Materials and Methods). In total, 2363 interacting proteins were identified (**Supplemental Table 3**), of which 1384 proteins were detected in both replicates (**Figure 1D**), and 1302 proteins fulfilled the consecutive binding criterion and were defined as C/EBPβ core interactions (βCI; **Figure 1E**).

### Validation of the PRISMA-derived dataset

To assess the biological significance of data derived by PRISMA, we compared PRISMA data (SET1, SET2, βCI) to previously identified C/EBPβ interactome data, as shown in **Figure 1F** (Siersbaek et al., 2011; Steinberg et al., 2012). In addition, we performed pull-down experiments using full-length C/EBPβ derived from SU-DHL1 cells and analysed the interactome by mass spectrometry (**Supplemental Table 4**). In total, 1369 proteins were significantly enriched in SU-DHL1 C/EBPβ samples compared with control samples using a false discovery rate (FDR) cutoff of 5%. The list of SU-DHL1 C/EBPβ interactors was included to extend the existing CEBPβ interaction datasets. The PRISMA core interaction data covered between 38% and 59% of the affinity purification-based datasets (see supplemental information). Affinity purification-based datasets were then combined and the overlap with PRISMA data was determined on the basis of their UniProt identifier entries, as shown in **Figure 1G** (see Materials and Methods). In total, 47% of data in all three sets were also found in the PRISMA core interactions, and 64% of PRISMA core interactions were also found in at least one of the other datasets. To estimate the FDR of the C/EBPβ interactor data detected by PRISMA, we employed the method proposed previously (D’Haeseleer and Church, 2004), which relies on comparing the intersections of protein interaction datasets to approximate the number of false-positive PPIs. We obtained FDRs of 11.2% and 13.9% for proteins detected in PRISMA replicates 1 and 2, respectively (SET1 and SET2 in **Figure 1D and E**; see Supplementary Information for details). FDRs were reduced to values below 4% when applying the filtering step leading to PRISMA core interactions. These results suggest that PRISMA data depict strong overlap and extensive coverage of the interactome related to native C/EBPβ. We conclude that the PRISMA method successfully extends the interactome data and serves as a resource for locating interaction footprints on C/EBPs.

### High-resolution C/EBPβ interactome footprints

The global protein interaction profile obtained from the C/EBPβ peptide matrix is depicted as a non-hierarchically clustered heat map in **Figure 2A**. The numerical distribution of proteins identified by individual peptides is shown in the upper part of **Figure 2A**. Peptides representing the DNA binding region (DB) and the C-terminal part of the leucine zipper (LZ) exhibit the highest number of protein interactions, yet locally-enriched binding hot-spots were also found with peptides from the N-terminal part of C/EBPβ.

**Figure 2:**
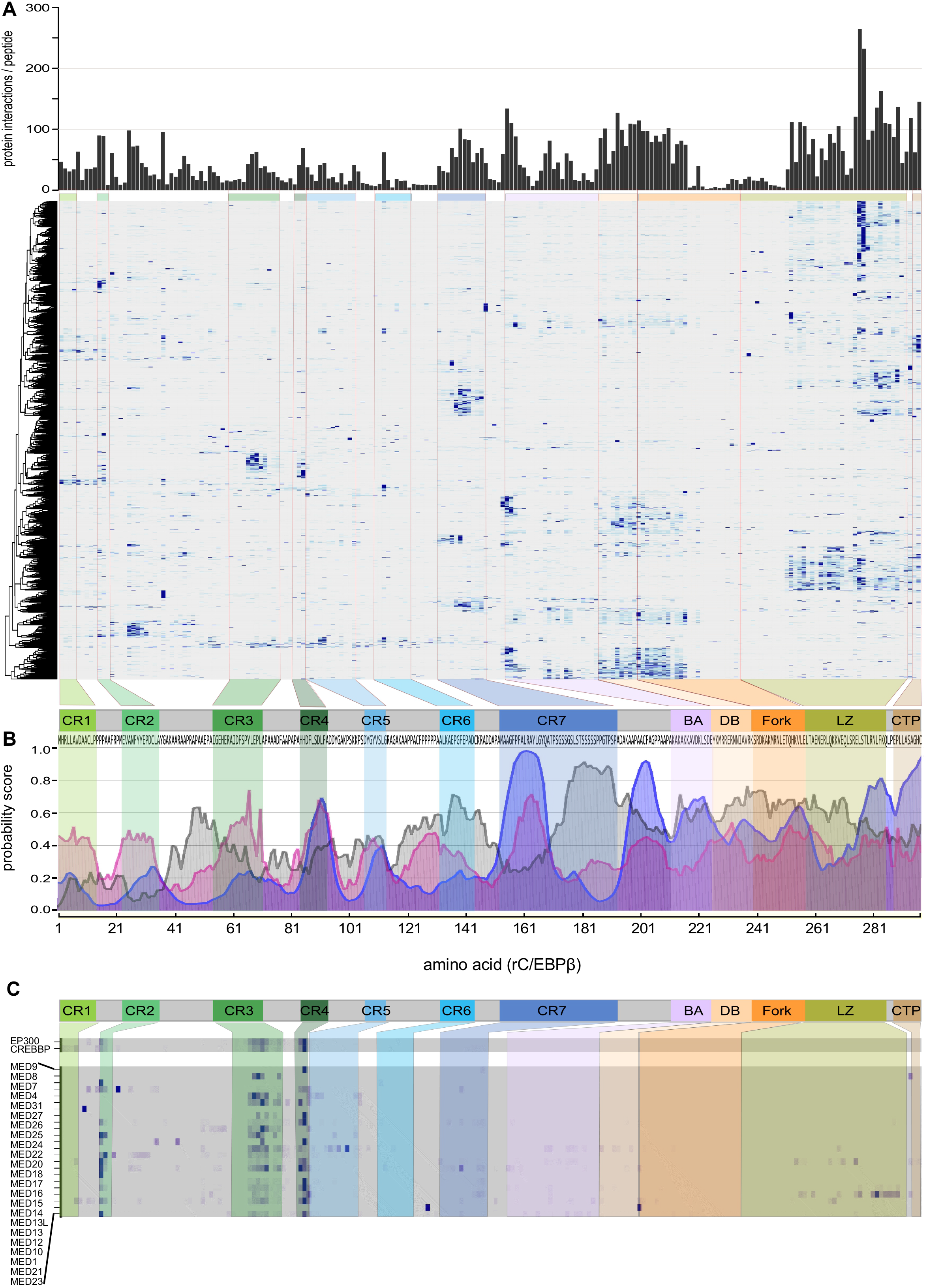
Distribution of protein interactions discovered by PRISMA. **A**. Heatmap of the normalised intensities of interaction partners. The bar-graph (top) shows the distribution of accumulated protein interactions by normalised binding intensities for each peptide. The tiled peptides of C/EBPβ (below) are plotted on the horizontal axis, and proteins are shown on the vertical axis following non-supervised hierarchical clustering. **B**. Prediction of C/EBPβ interaction regions using ‘Anchor’ (blue) and ‘MoRF’ (magenta) bioinformatics tools and prediction of intrinsic disorder (grey) in relation to the schematic representation of C/EBPβ and the primary sequence (bottom of A). Different conserved regions are coloured as shown in Figure 1A. **C**. Heatmaps of selected complexes.

Clusters of protein-peptide interactions tended to colocalise with regions predicted to undergo disorder-to-order transition during interaction with binding partners and coincide with the profile of conserved C/EBPβ regions (CR1–7), interaction and MoRF predictions (Disfani et al., 2012; Meszaros et al., 2009), as shown in **Figure 2B**. Predicted intrinsically disordered regions in the transactivating domain (CR1–4), IDR2 (between CR2 and CR3), and in the regulatory domain between CR7 and the basic/acidic region (IDR7) also displayed large numbers of interaction partners.

Next, PRISMA-derived data were compared with previously mapped C/EBPβ-interacting proteins. As shown in **Figure 2C**, both co-activator acetyltransferases, CBP and P300 (KAT3A/KAT3B) generated highly similar interaction footprints in the transactivation regions CR4 (strongest binding) and CR3 (additional binding), with some residual binding in CR2. Both C/EBPβ and C/EBPδ have been shown to interact with CBP/p300, and the main interaction region encompasses the CBP Taz2 domain as well as CR3 and CR4 in both C/EBPs. Removal of CR3 or CR4, or mutation of a critical tyrosine residue in CR3 or the DLF motif in CR4 all abrogated interaction with Taz2 and subsequent transcriptional coactivation (Kovacs et al., 2003; Schwartz et al., 2003). Previously published crystallographic data revealed that CR3/CR4 regions in C/EBPε may adopt an L-shaped α-helical structure that folds into the p300 Taz2 domain (Bhaumik et al., 2014), in agreement with the CBP/P300 protein footprints revealed by PRISMA (**Figure 2C**).

The multi-subunit Mediator (MED) complex (Conaway and Conaway, 2013; Jeronimo and Robert, 2017) has also been reported to interact with C/EBPβ (Mo et al., 2004), and several MED components were identified in a previously published C/EBPβ interaction dataset (Siersbaek et al., 2014) as well as in the SU-DHL1 interactome presented here. As shown in **Figure 2C**, PRISMA revealed 20 components of the multi-subunit Mediator (MED) complex that predominantly interact with CR2-, CR3-, and CR4-derived peptides from C/EBPβ TAD.

CR1 of the LAP* C/EBPβ isoform was reported to specify conjugation by SUMO3 (Eaton and Sealy, 2003) and in accordance, PRISMA showed interaction between SUMO3 and CR1. CR1 also interacts with the Brg1/SMARCA4 ATPase of the SWI/SNF/BAF complex. Moreover, the Brg1-CR1 interaction is sensitive to methylation of arginine 3 (Kowenz-Leutz et al., 2010). Consistent with these results, PRISMA revealed interaction of Brg1 with the unmodified CR1 peptide but not with the methylated peptide (**Supplemental Table 3**). Moreover, interactions with 10 additional protein components of BAF-SWI/SNF type protein complexes were detected with various peptides of the C/EBPβ transactivation domain and the bZip domain, indicating multiple interactions between C/EBPβ and chromatin modifying complexes, as previously suggested (Kowenz-Leutz and Leutz, 1999).

### The landscape of CEBPβ protein interactions

PRISMA data (SET1 and SET2) were inspected for enrichment of gene ontology (GO) terms, protein families and protein domains. As shown in **Table 1**, sorting of C/EBPβ interaction hits based on the quantity of terms resulted in exceedingly low Benjamini-Hochberg corrected FDRs (cut-off at a Benjamini-Hochberg corrected p-value of 0.01) (Finn et al., 2017; Finn et al., 2016; Gene Ontology, 2015). Strongly enriched terms included ‘nucleic acid binding’ (745 hits), ‘gene expression’ (611 hits), ‘protein binding’ (534 hits), ‘regulation of transcription’ (301 hits), ‘cell cycle’ (211 hits), ‘RNA splicing’ (172 hits), and several terms involving transcription, chromatin binding and remodelling. Gene set enrichment analysis showed that approximately 25% of PRISMA replicate validated proteins (267) fell into categories that involve GO terms related to RNA processing (FDR 5.63e-128). RNA binding proteins that contain a RNP-1 RNA binding domain fall into three classes that preferentially interact with the C/EBPβ bZip domain and the adjacent Fork and BA motifs, while a third group interacts with the CR2 region. A similar binding pattern was found for DEAD-box helicases, a group of enzymes involved in ATP-driven conformational adjustment of ribonucleoprotein assembly.

Protein domain enrichment using InterPro and PFAM databases revealed nucleotide binding proteins (102 hits), RNA binding proteins (80 hits), armadillo, WD40, histone fold, and DEAD box domain proteins as major C/EBPβ interaction partners. The number of domain hits obtained using standard algorithms to search protein domain databases may be underrepresented, because manual curation of PRISMA data increased the number of hits for WD40 domain proteins (IPR017986: WD40 repeat, region) from 32 to 40, and the number for DEAD box proteins (IPR011545: DNA/RNA helicase; DEAD/DEAH box type, N-terminal) from 20 to 24.

The C/EBPβ structure is predicted to fold back onto itself, permitting intramolecular signalling, or to stretch out to allow contact with several interaction partners simultaneously (Kowenz-Leutz et al., 1994; Lee et al., 2010a; Lynch et al., 2011). A strong indication of the involvement of multiple contacts with interaction partners through adjacent or distant C/EBPβ regions is shown in **Supplemental Figure 2A** that lists multivalent interactions, including potential interactions with different proteins of the same complex.

Discrete parts of the C/EBPβ primary structure have previously been assigned to different biochemical or cellular functions. Protein binding data from different C/EBPβ regions was extracted and GO term analysis performed for each of the regions to map structure-function relationships, as shown in **Supplemental Figure 2B**. While some functional attributions displayed partition over several regions of C/EBPβ (regulation of gene expression, RNA splicing), others are more localised to distinct regions. For example, basic and DNA binding regions shows strong enrichment for ‘regulation of mitotic cell cycle processes’ or ‘nuclear export’ whereas ‘nuclear import’ is associated with CR5, CR7, IR7 and LZ. While the C-terminal region of C/EBPβ is involved in the majority of GO terms, transcription factor coactivator functions are mainly associated with N-terminal activation of CR2, CR3, CR4 and IR7.

### C/EBPβ-interacting protein complexes

Next, PRISMA data were analysed for potential enrichment of large protein complexes interacting with CEBPβ. In addition to anticipated C/EBPβ interacting complexes such as the basic RNA polymerase II transcription machinery or Mediator, previously unknown interrelations became apparent. These included potential complexes involved in the RNA transcript processing machinery responsible for transcript capping, splicing, termination, and polyadenylation. Interactions included 3’end processing, cleavage and polyadenylating factor (CPSF1, 3, 7, 30, and 100), and several components of the Integrator complex involved in snRNA expression and the RNA exosome (EXOSC2, 4, 6, 7, 8, 9, and 10). Furthermore, many components of the nuclear pore and associated adapter complexes were identified. In addition, components of the transcript export THO-TREX complex, including THO1, 2, 3, 5, 6, 7, Aly, DDX39, and the THO-TREX-associated mRNA export factors Nxf1-Nxt1 were identified, along with a large number of heterogeneous nuclear ribonucleoproteins that may couple transcript splicing, maturation, and the formation of messenger ribonucleoprotein particles. Taken together, these data imply the existence of a previously unknown connection between CEBPβ and several steps involved in transcript generation, processing, maturation, and transport (Muller-McNicoll and Neugebauer, 2013; Wickramasinghe and Laskey, 2015).

Proteins co-occurring in both the PRISMA and SU-DHL1 proteomic data were then systematically explored to identify soluble protein complexes listed in the CORUM database (2017-03-06) using the g:Profiler bioinformatics toolkit (Reimand et al., 2016; Ruepp et al., 2010). A list of 1432 human protein complexes built from 2678 proteins (single UniProt identifier; ID) was derived after removal of redundant and non-human complex entries. Proteins listed in PRISMA replicates (SET1+SET2) and protein interactions derived from SU-DHL1 IP-mass spectrometry data were matched by UniProt and gene name ID using Perseus version 1.5.2.4 (Tyanova et al., 2016). Redundant gene names and isoform entries were merged into a single UniProt ID for each protein. Altogether, 816 proteins of the PRISMA-derived dataset and 490 proteins of the IP SU-DHL1 dataset were included in any of the 1432 CORUM complexes. The 1432 CORUM complexes were then ranked by a combination of two criteria: (i) the percentage of proteins in complexes obtained by PRISMA, and (ii) deviation from randomness of the coverage of the complexes obtained by PRISMA and SU-DHL1 (p-values with CORUM background, see Supplementary Information for details). Considering only complexes sharing at least one protein with the PRISMA dataset and which have at least three overlaps with PRISMA and SU-DHL1 data resulted in a ranked list of 417 candidate complexes (**Supplemental Table 5 and Supplemental Figure 3**). Whenever possible, complexes were grouped into categories indicating functional connections, such as DNA repair, nuclear pore or centromere, in addition to well-characterised multi-subunit complexes, such as Mediator, SWI/SNF, MLL, NuRD, and others. As shown in **Figure 3A**, 45 multi-protein complexes and categories were extracted, and normalised protein interaction values at any of the 203 C/EBPβ peptides were summed over all instances of the category in the list and depicted as a bar-plot to visualise the distribution of interaction sites of different categories or complexes in relation to the C/EBPβ primary sequence. These 45 entities were composed of 30 complexes or categories representing the full upper quartile of the ranked complex list (i.e. up to rank 104, **Supplemental Table 5**, with minor exceptions, **Supplemental Figure 3**), together with 15 lower-ranking functional counterparts. **Figure 3B** shows prominent node-link diagrams of a selection of 14 multi-protein complexes among the highest-ranking categories that share many proteins identified by both PRISMA and SU-DHL1 immunoprecipitation methods.

**Figure 3:**
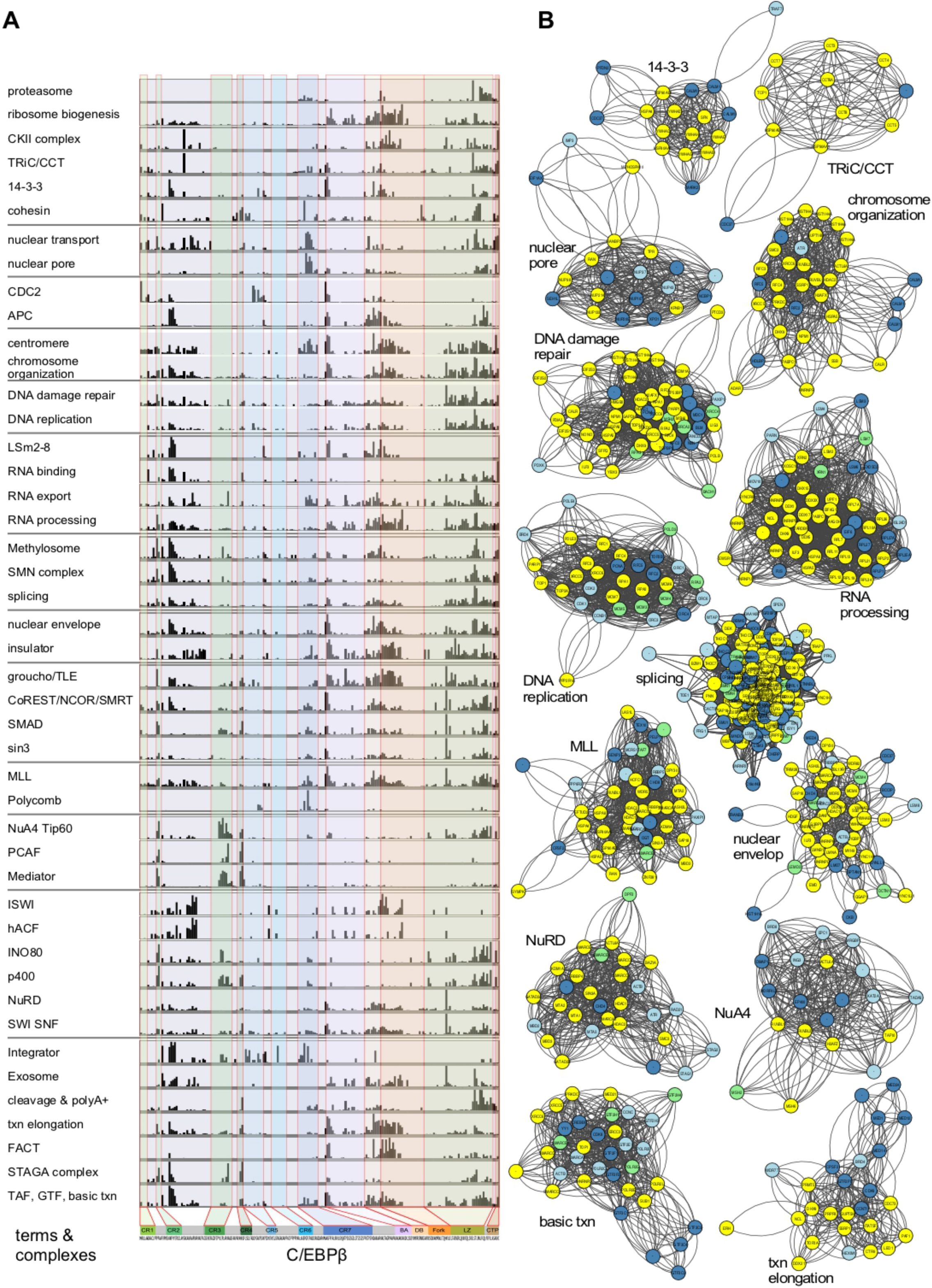
PRISMA-based prediction of protein complexes interacting with C/EBPβ. **A**. Bar-graphs of accumulated normalised intensities for 45 selected complexes. Based on CORUM database protein complex annotations, the corresponding normalised intensities of proteins identified by PRISMA were extracted and plotted. **B**. Network representation of 14 potential complexes identified. Nodes are colour-coded according to their detection in PRISMA and SU-DHL1 IP experiments (yellow = detected in IP and PRISMA, green = detected in IP, dark blue = detected in the PRISMA core set, light blue = detected in PRISMA SET1 and 2).

### Validation of region- and PTM-specific C/EBPβ interactions

Next, conventional protein pull-down, immunoprecipitation, and immunoblotting analysis was used in combination with CEBPβ mutants to validate data from the PRISMA-derived interactome (**Figure 4**). According to PRISMA, RelA and NUP50 both predominantly interacted with peptides located in CR7. Bacterially expressed GST-C/EBPβ constructs were probed with HEK293 extracts to examine RelA and NUP50 interactions. As shown in **Figure 4B**, affinity capture with GST-CEBPβ constructs and immunoblotting confirmed CR7 as the interaction site for both proteins.

**Figure 4:**
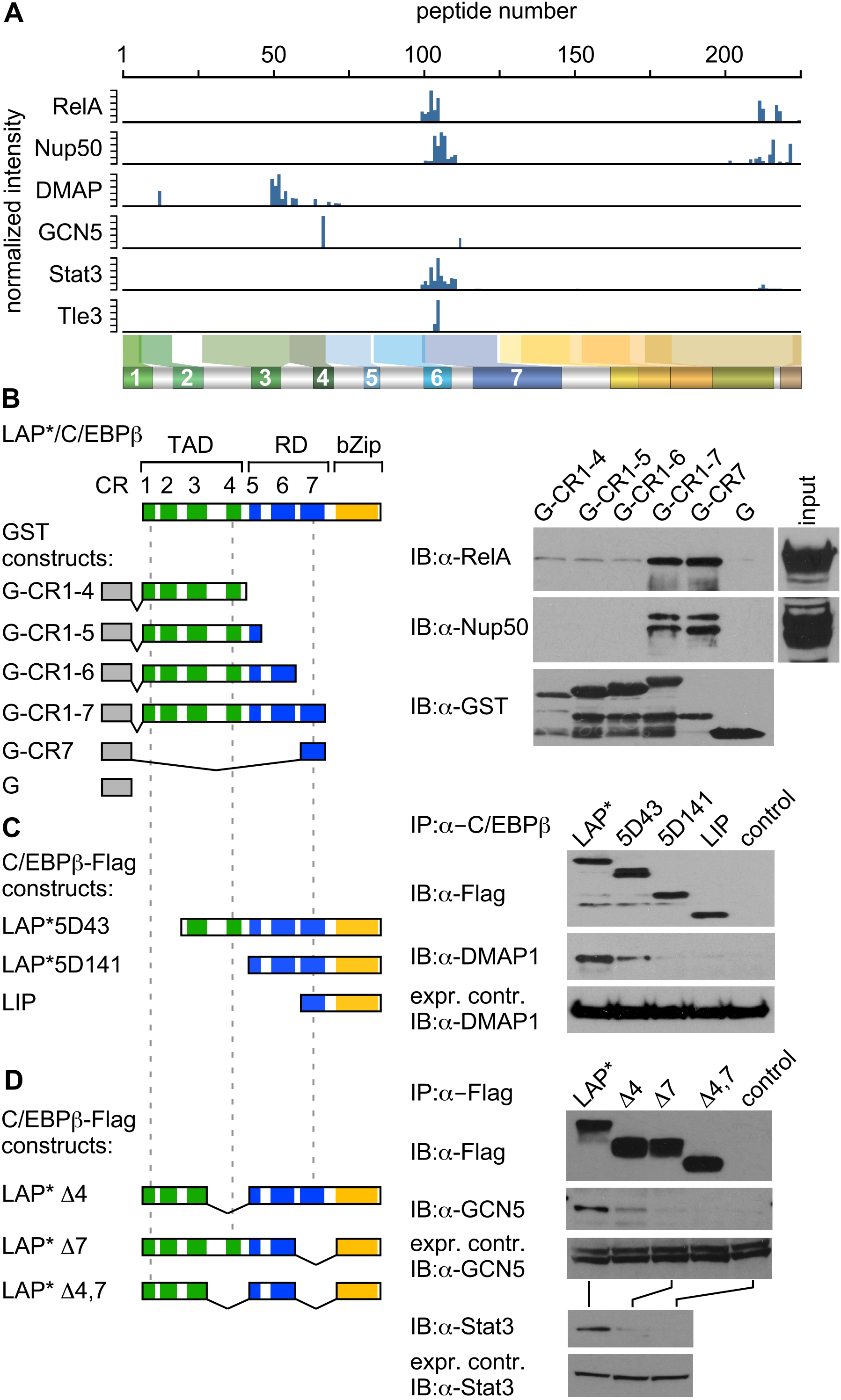
Validation of site-specific interactions with C/EBPβ. **A**. Accumulated binding intensities of interaction partners (indicated on the left) from the PRISMA screen. Bar-graphs indicate interactions according to the peptide position in C/EBPβ, as schematically shown underneath. **B**. Top: Colour-coded scheme of the C/EBPβ protein with dashed lines to aid comparison of constructs shown in B, C, and D. Bottom: Bacterially expressed GST-C/EBPβ constructs (left) and immunoblots (right) showing interaction of RelA and Nup50 in nuclear cell extracts with the different C/EBP β deletion mutants. The most intense signals of RelA and Nup50 are associated with the presence of the CR7 region of C/EBPβ. A, Antibody; IP, immunoprecipitation; IB, Immunoblotting. **C**. Co-immunoprecipitation of FLAG-tagged LAP* C/EBPβ or three N-terminal C/EBPβ deletion mutants, followed by immunoblot detection of co-precipitated DMAP1. **D**. Immunoprecipitation of LAP* and three different internal deletion mutants of C/EBPβ followed by immunoblot detection of GCN5 and Stat3.

The histone acetyltransferase GCN5 of the SAGA complex and the DNA methyltransferase-associated protein DMAP1 of the NuA4/TIP60 complex are involved in a wide variety of developmental and genome regulatory activities, including transcription, DNA repair, DNA methylation, and chromatin remodelling (Mohan et al., 2006; Weake and Workman, 2012). DMAP1 and GCN5 engage in major binding to CR3 and CR4, respectively, and minor binding elsewhere in C/EBPβ. Various C-terminally FLAG-tagged C/EBPβ mutant constructs lacking CR1, CR2 (5D43), the entire TAD (5D141), CR4 (LAP*Δ4), CR7 (LAP*Δ7), or both (LAP*Δ4,7) were expressed in HEK293 cells to examine selective and multi-site interaction with resident DMAP1/GCN5 or their respective complexes. As shown in **Figure 4C and D**, pull-down and immunoblotting revealed that removal of the DMAP1 minor binding site in CR2 partially abrogated the interaction, and removal of binding to CR3 completely abrogated the interaction. Similarly, removal of the major binding site for GCN5 located in CR4 strongly affected GCN5 association, and deletion of the ancillary binding site in CR7 entirely abolished binding to C/EBPβ.

### The impact of PTMs on C/EBPβ-mediated interactions

Detection of PTM-dependent protein interactions in linear C/EBPβ peptides was a major objective for the development of the PRISMA screening method. Protein binding to each PRISMA peptide and its PTM-modified versions were therefore examined. The binding signal for each protein was normalised against the signal from the corresponding unmodified peptide to compare enhanced or reduced binding. Interacting proteins were then clustered, as shown in **Supplemental Figure 4**. Depending on PTM, as exemplified in **Figure 5A**, interactions fell into four categories; (i) PTM-independent binding, (ii) generally enhanced binding, (iii) generally repressed binding, and (iv) PTM-specific binding. Most of the identified binding partners were indifferent to the type of PTM and were classified as binding proteins that may recognise parts of the primary peptide sequences. Generally enhanced binding may in part relate to the fact that many PTMs increase hydrophobicity and may thus enhance interactions non-specifically. The most interesting binding partners are expected to be found in the last two categories and represent proteins that are disturbed by any PTM in the peptide or represent interaction partners that were attracted or repelled by a particular PTM. Examples of the latter are DMAP1 binding to CR3 or RelA, and TLE3 binding to CR7, the binding of which depends on the methylation status of arginine residues that were subsequently examined.

**Figure 5:**
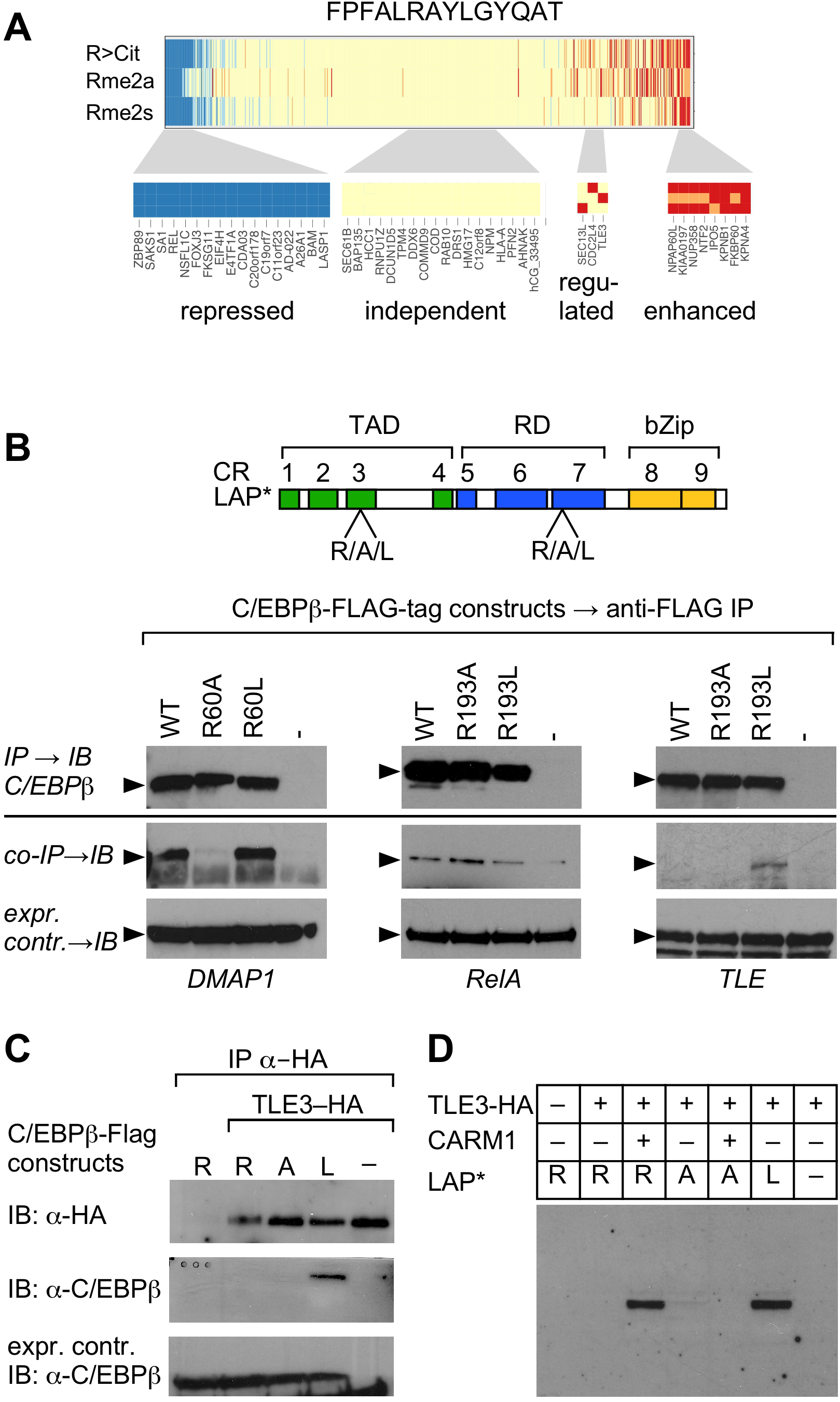
PTM-specific interactions with C/EBPβ. **A**. PTM-dependent binding to the FPFALRAYLGYQAT peptide of C/EBPβ CR7. Normalised binding intensity is relative to the unmodified peptide. Binding can be categorised into four different types; proteins not responding to modifications on C/EBPβ (yellow, middle), proteins with reduced binding independent of the PTM (blue, left), proteins with enhanced binding independent of the nature of the PTM (red, right), and specific PTM-dependent binding (yellow-red, small). **B**. Co-immunoprecipitation with wild-type (WT) C/EBPβ and two mutants abrogating (R→A) or mimicking (R→L) methylation at R60 or R193. Binding of DMAP1 was followed by immunoblotting. DMAP is able to bind to WT and R→L mutant proteins, but replacement of R60 with an alanine suppresses the binding of DMAP. Similarly, TLE3 shows an enhanced binding to the R→L mutant compared with the WT protein the R→A mutant. RelA shows a slightly different binding behaviour, preferring the alanine mimic over the leucine mutant and WT forms. B. Inverse immunoprecipitation experiment with an HA-tagged version of TLE3 was co-expressed with C/EBPβ and two mutants carrying the arginine methylation mimics R193A and R193L. Detection of C/EBPβ shows enhanced binding of C/EBPβ to R193L C/EBPβ. C. Co-expression of full-length C/EBPβ (LAP*), TLE3 and the histone-arginine methyltransferase CARM1. TLE3 in combination with CARM1 can precipitate WT C/EBPβ, similar to the arginine-methylation mimic R193L.

The transactivation region CR3 (residues 53–68: AIGEHERAIDFSPYLE) contains an arginine residue that is conserved in C/EBPβ but not in C/EBPα,δ,ε and was previously found to be methylated (Leutz et al., 2011). The PRISMA data suggested that DMAP1 binding to CR3 depends on methylation of R59. Another interaction hot-spot was mapped to the start site of the highly conserved, alternatively initiated LIP C/EBPβ isoform that also represents part of the regulatory region CR7 (residues 158–170: FPFALRAYLGYQAT). The respective arginine residues in both chicken (R60, R193) and rat (R58, R162) were previously found to be methylated. Mutation of the equivalent amino acid residue to alanine/leucine in chicken C/EBPβ strongly altered transcriptional activity, suggesting that the methylation status of the conserved arginine in CR7 may be critical (Leutz et al., 2011). WT C/EBPβ, methylation-defective R60A and R193A, and methylation mimicking R60L and R193L constructs were subjected to co-immunoprecipitation to compare alterations in binding, as shown in **Figure 5B**. As suggested by PRISMA, interaction of all three proteins with C/EBPβ was sensitive to the amino acid side chain configuration at the respective arginine positions, demonstrating both amino acid and PTM specificity, in addition to evolutionary conservation of chicken and rat C/EBPβ interactions. Binding of DMAP1 to C/EBPβ tolerated the R60L exchange but not the R60A exchange, confirming the side chain specificity and preference for increased side chain hydrophobicity. RelA bound more strongly to the R193A mutant than the R193L mutant, confirming rejection by increased hydrophobicity, as represented by methylation of R193. By contrast, the TLE3 co-repressor (Agarwal et al., 2015) strongly favoured binding to the R193L mutant, suggesting an increase in hydrophobicity but not the positive charge of the arginine side chain was important for interaction with TLE3. The immunoprecipitation order of TLE3 was thus reversed to further validate the preference of TLE3 for the R to L mutation. As shown in **Figure 5C**, co-immunoprecipitation of C/EBPβ with TLE3 preferentially occurred with C/EBPβ R193L, but not with R193 or R193A. Next, we identified R193 in C/EBPβ as a target of the arginine methyltransferase PRMT4/CARM1. As shown in **Figure 5D**, co-immunoprecipitation of TLE3 together with WT C/EBPβ was dependent on co-expression of PRMT4/CARM1. Importantly, and irrespective of the presence of PRMT4/CARM1, the methylation-defective C/EBPβ R193A mutant failed to bind TLE3, confirming C/EBPβ CR7 arginine side chain methylation as a prerequisite for binding of the TLE3 groucho-type co-repressor. We conclude that many C/EBPβ interactions, as exemplified by DMAP1, RelA, and TLE3, are methylation sensitive and that PRISMA was able to uncover PTM-mediated regulation of PPIs.

### Novel connections between C/EBPβ, transcription elongation, MLL, and NuRD

Of particular interest was the observation that many proteins forming part of the machinery involved in RNA polymerase II (RNAPII) pausing and elongation were included in the PRISMA data. Components of the trithorax MLL/Set1/COMPAS histone H3 lysine 4 (H3K4) methyltransferase complexes (Shilatifard, 2012) were also represented in the dataset, raising the possibility of a connection between C/EBPβ, enhancer/promoter binding, and regulation of RNAPII processivity. Many components implicated in both processes were found, including the MLL complex components CHD3 and 4 (Mi-2), ASH2, Dpy30, PTIP, HCF-1, the general transcriptional elongation factor B (TFIIS) and associated elongin A/B/C factors (Aso et al., 1995), the P-TEFb components cyclinK/T1, cdk9, and additional regulatory components including LARP7, HEXIM1, SPT5 (DSIF), and the chromatin adaptor bromodomain factor Brd4 and components of the super elongation complex (SEC) including AF9, PCAF1, and AF4 (AFF1) (Luo et al., 2012). AFF1 is a central SEC scaffold component, and AF9 and ENL are highly similar YEATS domain family members that compete for binding to AFF1. The N-terminal YEATS domain of AF9 and ENL both bind to the PAF complex to connect SEC to RNAPII on chromatin templates. The CDK9/CYCT1 P-TEFb complex is required for rapid transcriptional induction, phosphorylation of the C-terminal domain of RNAPII, and engagement of BRD4, which also interacts with H3K9ac. P-TEFb is also associated with a 7SK snRNA subcomplex that contains the regulatory components LARP7, HEXIM1, and SPT5 (DSIF) and connects to PAFc in the dynamic transcription elongation machinery (He et al., 2011; Luo et al., 2012; Peterlin and Price, 2006). In addition, the histone chaperone FACT complex facilitates passage of the transcription apparatus through chromatin and is thought to restore the chromatin structure and the epigenetic state during transcription, replication and repair (Hammond et al., 2017; Hondele and Ladurner, 2013; Orphanides et al., 1998).

Immunoprecipitation and immunoblotting was performed in order to validate several of these novel connections. As shown in **Figure 6**, multiple MLL, FACT, and SEC components were all co-immunoprecipitated with C/EBPβ, confirming the predictive capacity of PRISMA. It is also important to note that AFF1, AF9, and ENL are among the most frequent oncogenic fusion partners with the MLL gene product that transforms early haematopoietic progenitors and causes childhood leukaemia by short-circuiting enhancer and promoter activation and transcriptional elongation checkpoint controls (Krivtsov and Armstrong, 2007; Shilatifard, 2012; Slany, 2009; Smith et al., 2011).

**Figure 6:**
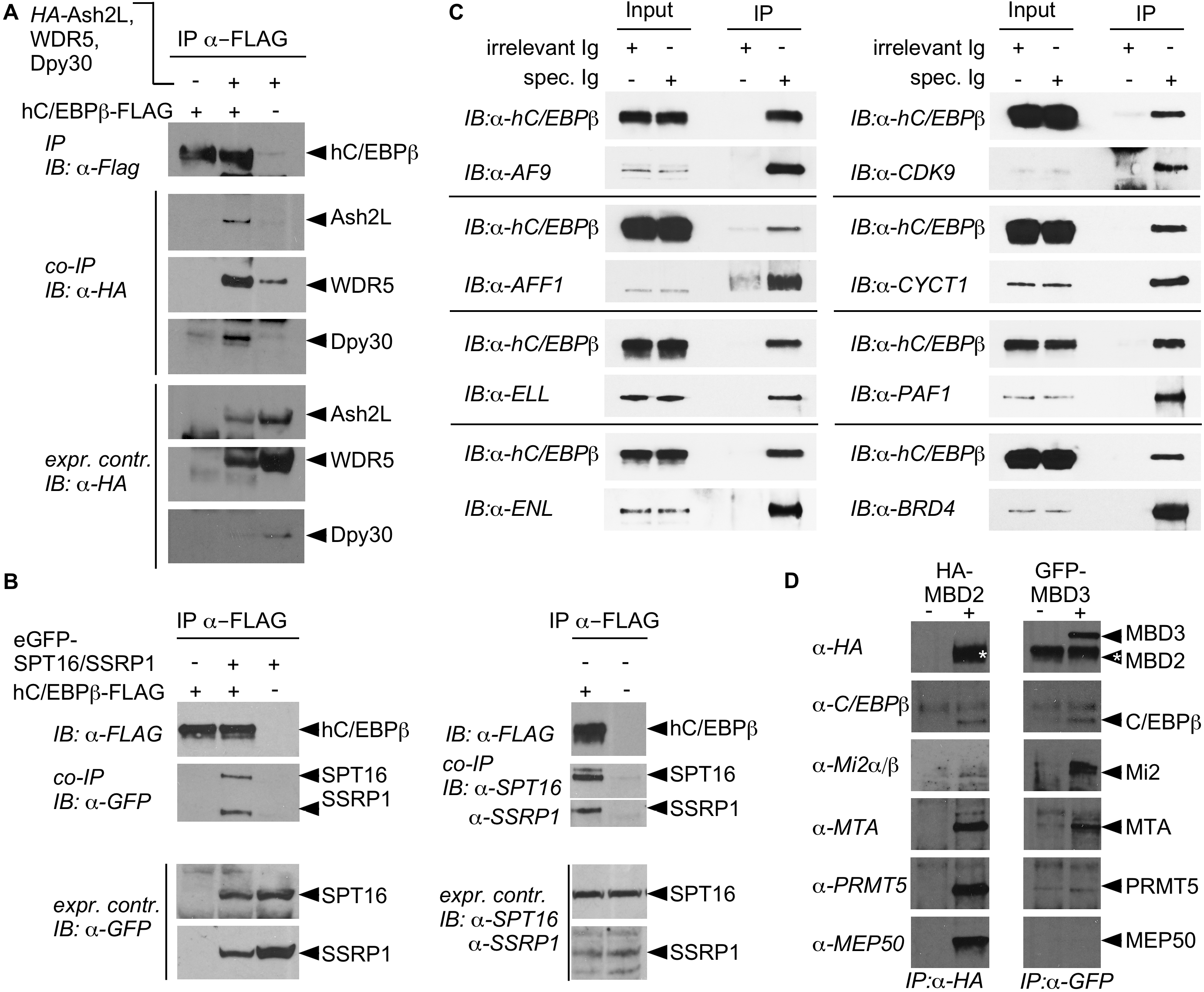
C/EBPβ-interacting multi-protein complexes. **A**. Co-immunoprecipitation of FLAG-tagged C/EBPβ and the Ash2/Wdr5 complex. **B**. FLAG-tagged C/EBPβ was co-expressed with GFP-tagged Spt16 and SSRP1. Anti-Flag coprecipitates both FACT complex subunits. **C**. Interaction with the super elongation complex. Immunoprecipitation of human C/EBPβ (C/EBPβ) pulls down the entire super-elongation complex. **D**. Interaction with the NuRD complex. The integral subunits of the NuRD complex MBD2 (HA-tagged) and MBD3 (GFDP-tagged) were pulled down by immunoprecipitation. Detection by western blotting shows that C/EBPβ interacts with both subunits.

The Mi2/**Nu**cleosome **R**emodelling **D**eacetylase (NuRD) corepressor complex is widely expressed and maintains chromatin in a closed state via ATP-dependent chromatin remodelling and histone tail deacetylation. Some of the NuRD components (CHD3/4(Mi2), HDAC1, RBBP4/7, MTA1,2, GATAD2A/p66α, MBD2, MBD3) identified by PRISMA, SU-DHL1 IP and other approaches (Steinberg et al 2012; Siersbaek et al 2014) are shared with the SIN3A and CoREST repression complexes (Gregoretti et al 2004), and all were implicated by CORUM analysis (**Figure 3, Supplemental Figure 3, Supplemental Table 4**). Two major NuRD complexes contain either the unique methyl-CpG binding proteins MBD3 or MBD2, the latter of which may also associate with the WDR77(MEP50)/PRMT5 sub-module (Le Guezennec et al., 2006; Zhang and Yinghua, 2011). Immunoprecipitation of either MBD2 or MBD3 revealed association of C/EBPβ, Mi2, and MTA with both MBD components, whereas only MBD2 co-precipitated with the WDR77(MEP50)/PRMT5 sub-module, suggesting that C/EBPβ may associate with both types of NuRD complexes.

In conclusion, the PRISMA technique reveals a comprehensive and PTM-dependent interactome of the disordered hub transcription factor C/EBPβ, and permits the detection of footprints of protein interactions and protein complexes. It appears that C/EBPβ is involved at the nexus between enhancer and promoter regulation, in all aspects of pre- and posttranscriptional control including initiation and pausing control, splicing and exosomal RNA degradation, polyadenylation, RNA maturation and nuclear export. PRISMA has been developed for analysis of the interactome of proteins with a high degree of intrinsic disorder. Such proteins lack a defined 3D structure, and protein interactions and complex formation are based on SLiMs, PTMs, and intrinsic flexibility. SLiM interactions often occur with high specificity but low affinity to enable multiple contacts of a dynamic nature and IDRs in conjunction with PTMs may facilitate structural transitions in partner protein exchanges (Wright and Dyson, 2015). The high local concentration of peptides in matrix spots may limit free diffusion and thus overcome the affinity/dissociation problem of low affinity interactions (Ruthenburg et al., 2007). These features permit the detection of weak interactions and interactions with rare protein partners. The interaction screen developed herein is augmented by inclusion of amino acids with modified side chains to enable detection of PTM-dependent interactions on a global scale. Together, the footprints of interactions and PTM-dependent data may help in the rational design of mutants to explore the functional C/EBPβ interactome, and aid in the development of screens for pharmacological inhibitors of interactions.

## Material & Methods

### Peptide matrix synthesis

Peptides were synthesised using the automated high-throughput SPOT-synthesis method (Kramer and Schneider-Mergener, 1998). Briefly, Whatman 50 cellulose membranes (Whatman, UK) were functionalised by coupling of Fmoc-protected β-alanine in defined spots. Subsequently, peptides were synthesised stepwise using standard Fmoc-chemistry. After each coupling step before Fmoc-deprotection, peptides that failed coupling of building blocks were acetylated to avoid false sequences.

### Interaction screen

The peptide matrix membrane was blocked with yeast tRNA (1 mg/ml; Invitrogen, Karlsruhe, Germany) in binding buffer (20 mM HEPES pH 7.9, Merck, Germany, 0.2 mM EDTA, Merck, Germany, 100 mM KCl Merck, Germany, 20% glycerol, Merck, Germany, 0.5 mM DTT, Merck, Germany) to minimise unspecific protein binding. The membrane was then incubated with HeLa cell nuclear extract (5 mg/ml; Calbiotech S.A, Germany) in binding buffer supplemented with 0.5 mM PMSF (Merck, Germany) for 30 min, then briefly washed with binding buffer. Peptide spots were individually excised and bound proteins converted to peptides using a two-step digest with endopeptidase LysC (Wako, Japan) followed by sequencing-grade trypsin (Promega, Germany) using a robotic setup (Kanashova et al., 2015). Peptide extracts were purified and stored on stage tips (Rappsilber et al., 2007). Two replicates were measured for each peptide spot.

### Mass spectrometry measurement for PRISMA

Peptides were cleaned up using a stage-tip micro column (Rappsilber et al., 2007) and resuspended in water with 0.1% formic acid (Merck, Germany). Peptides were separated on a 15cm reverse-phase column (packed in-house, 75 μm inner diameter, 3 μm C18-Reprosil beads; Dr Maisch GmbH, Ammerbruch, Germany) using a gradient to 40% acetonitrile (Merck, Germany) developed over 1 h 5 min. Separated peptides were ionised on a Proxeon ion source and directly sprayed into the online-coupled VELOS-OrbiTRAP mass spectrometer (Thermo scientific). MS^1^ spectra were recorded with a mass resolution of 60,000 in the orbitrap part of the machine. MS^2^ spectra were recorded in the VELOS. The ten most intense ions with a charge state greater than 1 were selected (target value = 500; monoisotopic precursor selection enabled) and fragmented in the linear quadrupole trap using CID (collision induced dissociation, 35% normalised collision energy). Dynamic exclusion for selected precursor ions was 60 s. Recorded spectra were analysed using MaxQuant software package version 1.2.2.5 (Cox and Mann, 2008) and the human IPI database (version 3.3.72), allowing for 2 missed cleavages. Fixed modifications were set to cysteine carbamylation, and variable modifications were set to methionine oxidation, as well as N-terminal protein acetylation. Each replicate was analysed separately with the label-free option activated for data quantification (Cox et al., 2011).

### Analysis of nuclear cell extracts

Nuclear cell extracts were supplemented with the USP2 standard (Merck, Germany) and digested as described above on an automated digestion setup (Kanashova et al., 2015). Peptides were fractionated by RP-HPLC with Proxeon nLC2 and further analysed by a QExactive mass spectrometer (Thermo Scientific). The mass spectrometer was operated in a data-dependent acquisition mode with dynamic exclusion enabled (30 s). MS^1^ (mass range 300-1700 Th) was acquired at a resolution of 70,000 with the ten most abundant multiply charged (z ≥ 2), ions selected with a 2 Th isolation window for HCD (Higher-energy collisional dissociation) fragmentation. MS^2^ scans were acquired at a resolution of 17,500 and injection time of 60 ms.

### Full-length C/EBPβ interactome analysis by AP-MS mass spectrometry

Eluates from control immunoglobulins (IgG) and anti-C/EBPβ pull-downs (four replicates each) were ethanol precipitated and protein pellets were solubilised in urea buffer (6 M urea, 2 M thiourea, 20 mM HEPES pH 8), reduced for 30 min at RT in 10 mM DTT, followed by alkylation with 55 mM chloroacetamide (Merck, Germany) for 20 min in the dark at RT. The endopeptidase LysC (Wako, Japan) was added at a protein:enzyme ratio of 50:1 and incubated for 4h at RT. After dilution of the sample with 4× digestion buffer (50 mM ammonium bicarbonate pH 8), sequence-grade modified trypsin (Promega, Darmstadt, Germany) was added (protein:enzyme ratio = 100:1) and digested overnight. Trypsin and LysC activity was quenched by acidification with trifluoroacetic acid (TFA) added to pH ~2. Afterwards, peptides were extracted and desalted using the standard StageTip protocol (Rappsilber et al, 2003). Peptide mixtures were separated by reverse-phase chromatography using an Eksigent NanoLC 400 system (Sciex, Darmstadt, Germany) on in-house-manufactured 20 cm fritless silica microcolumns with an inner diameter of 75 μm. Columns were packed with ReproSil-Pur C18-AQ 3 μm resin (Dr Maisch GmbH). Peptides were separated using an 8–60% acetonitrile gradient (ran over 224 min) at a nanoflow rate of 250 nl/min. Eluting peptides were directly ionised by electrospray ionisation and analysed on a Thermo Orbitrap Fusion instrument (Q-OT-qIT, Thermo). Survey scans of peptide precursors from 300 to 1500 *m/z* were performed at 120 K resolution with a 2×10^5^ ion count target. Tandem MS was performed by isolation at 1.6 m/z with the quadrupole, HCD fragmentation with a normalised collision energy of 30, and rapid scan MS analysis in the ion trap. The MS^2^ ion count target was set to 2×10^3^ and the max injection time was 300 ms. Only precursors with a charge state of 2–7 were sampled for MS^2^. The dynamic exclusion duration was set to 60 s with a 10 ppm tolerance around the selected precursor and its isotopes. The instrument was run in top speed mode with 3 s cycles, meaning the instrument could continuously perform MS^2^ events until the list of nonexcluded precursors diminished to zero or 3 s. Data were analysed by MaxQuant software version 1.5.1.2. The internal Andromeda search engine was used to search MS^2^ spectra against a decoy human UniProt database (HUMAN.2014-10) containing forward and reverse sequences. The search included variable modifications of methionine oxidation and N-terminal acetylation, deamidation (N and Q) and fixed modification of carbamidomethyl cysteine. The minimal peptide length was set to seven amino acids, and a maximum of two missed cleavages were allowed. The FDR was set to 0.01 for peptide and protein identifications. Unique and razor peptides with a minimum ratio count of 1 were considered for quantification. Retention times were recalibrated based on the built-in nonlinear time-rescaling algorithm. MS^2^ identifications were transferred between runs with the ‘Match between runs’ option, in which the maximal retention time window was set to 0.7 min. Statistical analysis was performed using Perseus version 1.5.2.4. C/EBPβ pull-down and control samples were defined as groups and proteins and filtered by intensity value using a ‘minimum value of 3 per group’ as the threshold. After log2 transformation, missing values were imputed with random numbers from a normal distribution with a mean and standard deviation chosen to best simulate low abundance values below the noise level (width = 0.3; shift = 1.8). Significantly enriched proteins were determined using a volcano plot-based strategy, combining standard two-sample t-test *p*-values with ratio information. Significance corresponding to an FDR of 5% was determined by a permutation-based method (Tusher et al., 2001). Equal sample load was confirmed by calculating the ratio of antibody intensities (mean log2 ratio = 0.1575).

### Identification of C/EBPβ PTM sites by mass spectrometry

Eluates of anti-C/EBPβ pull-downs were ethanol precipitated and protein pellets were processed as described above for the interactome analyses. Peptides were analysed on a Thermo Orbitrap Fusion instrument (Q-OT-qIT, Thermo). Sequential survey scans of peptide precursors covering different mass ranges (300–600, 550–850, 800–1100, 1050–1700 m/z) were performed at 120 K resolution with a 2×10^5^ ion count target on the three most abundant precursor ions. Tandem MS was performed by isolation at 1.6 m/z with the quadrupole, HCD fragmentation with a normalised collision energy of 30, and rapid scan MS analysis in the ion trap. The MS^2^ ion count target was set to 1×10^4^ and the max injection time was 500 ms. Only precursors with a charge state of 2–7 were sampled for MS^2^. The dynamic exclusion duration was set to 60 s with a 10 ppm tolerance around the selected precursor and its isotopes. Data were analysed by MaxQuant software version 1.5.1.2 as described above for interactome analysis with some modifications; the search included variable modifications of methionine oxidation and N-terminal acetylation, deamidation (N and Q), phosphorylation (S, T and Y), acetylation (K), methylation (K and R), dimethylation (K and R), trimethylation (K) and citrullination (R). Modification of carbamidomethyl cysteine was set as a fixed modification. The minimal peptide length was set to seven amino acids, and a maximum of four missed cleavages were allowed. The FDR was set to 0.01 for site identifications. To filter for confidently identified peptides, the MaxQuant score was set to a minimum of 40. Identified PTM sites were classified according to their localisation probability (Class I >0.75, Class II >0.5, Class 3 >0.25). In order to distinguish citrullination from deamidation, a modification resulting in a similar mass shift of the precursor, both modifications were included during MaxQuant data analyses as variable modifications, and at least one missed cleavage was required for citrullination site identification. When arginine and lysine are acetylated or methylated, trypsin often fails to cleave at that site, resulting in miss-cleaved peptides. Therefore, only the PTM sites, which were identified within the peptide, but not C-terminally localised, were considered as confidently identified.

### Combining the two datasets and data filtering

The two datasets of the PRISMA measurement were analysed in two batches due to restrictions of the MaxQuant software package. The two separate datasets were then integrated by calculating the average intensity value where two values were available, or by taking the single measured value if only one intensity measurement was available to prevent bias against single identifications. Single value intensities were annotated as lower confidence quantifications. Each of the data rows of the combined dataset, corresponding to intensity values of one protein over all 203 peptides, was first filtered according to the outlier criterion *I_n_* = 0 *if I_n_* ≤ *P*90, where In is the intensity at peptide n, and P90 is the 90th percentile of the intensity value distribution of the protein, followed by normalisation against the intensity of the highest value in each row using 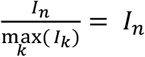, where I_n_ is the intensity at peptide n. This was followed by an additional filtering step where all proteins were removed that did not show a consecutive binding pattern according to finding at least one intensity I_n_ with I_n_ > 0 and I_n+1_ > 0 with I_n_ as the intensity at peptide n and only considering peptides without any PTM.

### Construction of CORUM networks

For each complex identified in the CORUM database, the corresponding UniProt identifier was extracted and translated to an ensemblp identifier using the bioDBnet conversion tool (Mudunuri et al., 2009). For each of the ensemblp identifiers, interactions were extracted from the STRING resource (Szklarczyk et al., 2015). The interaction network was constructed using the igraph software package (Csardi G, 2006) and coloured based on the source the of the interaction from the different datasets.

### Relative binding to PTM-modified peptides

For each peptide and its PTM derivatives, the relative binding was calculated. For each protein and modified peptide, the intensity value of the unmodified form of the peptide was subtracted. The resulting values were separated into five fractions and clustered according to their Euclidian distance.

### Calculation of peptide properties

For calculation of peptide properties, the R peptides package was used, and properties were calculated and plotted using the ggplot2 packages (Wickham, 2009).

### Use of the DAVID package

A total of 1375 identifiers were examined using the Generic Gene Ontology Term Finder (GGOTF, Princeton University, Lewis-Sigler Institute for Integrative Genomics) to assess ontology aspects, functions, and components.

### Immunoprecipitation of endogenous C/EBPβ for mass spectrometry

Immunoprecipitation of C/EBPβ from 4×10^8^ SU-DHL1 cells (anaplastic large cell lymphoma, DSMZ, ACC 356) was performed after washing twice with phosphate-buffered saline (PBS) and resuspension in lysis buffer (20 mM HEPES pH 7.5, 150 mM NaCl, 5 mM MgCl2, 1 mM EDTA pH 8), 1 μM ZnCh_2_ (Merck, Germany), 0.1% NP40 (Sigma, Germany), 2 mM dithiothreitol (DTT), 2 mM PEFAbloc (Böhringer, Mannheim, Germany) supplemented with protease inhibitor cocktail (Roche, Germany), and 20 U/ml benzonase (Sigma, Germany). After incubation on ice for 10 min, lysates were sonicated twice for 1 min, cell debris was removed by centrifugation at 70,000 g for 30 min, and lysates were filtered through a 0.45 μM filter (Whatman, Maidstone, UK) prior to immunoprecipitation. Samples were immunoprecipitated with a C/EBPβ antibody mix for 30 min at 4°C (Santa Cruz; C-19 and a customised polyclonal antibody raised against the human recombinant C/EBPβ protein) and immunoprecipitates were subsequently collected on Protein-G Dynabeads (Novex, Life Technologies). Beads were washed twice in lysis buffer without benzonase, once in lysis buffer without benzonase and NP40, and eluted by incubation with a mix of 6 M urea and 2 M thiourea (Sigma) for 15 min at 25°C. Immunoprecipitation specificities were controlled by immunoprecipitation of SU-DHL1 lysates with nonspecific rabbit IgG control antibodies (Santa Cruz, sc-2017) and subsequent collection on Protein-G Dynabeads.

### Cell culture, transfection, immunoprecipitation and immunoblotting

HEKT-293 cells were grown in Dulbecco’s modified Eagle medium (DMEM; Invitrogen, USA) and SU-DHL1 cells were grown in RPMI (Invitrogen, USA) supplemented with 10% FCS and 1% penicillin/streptomycin (Invitrogen, USA). Transfection of plasmids in HEKT-293 cells was performed by calcium-phosphate precipitation or Metafectene (Biontex, Munich, Germany) according to the manufacturer’s protocol. For validation of PRISMA-identified C/EBPβ protein interactions, immunoprecipitation of WT or mutant C/EBPβ proteins expressed in HEKT-293 cells was performed as described previously (Kowenz-Leutz et al, 2010). Briefly, cell lysates were prepared in lysis buffer and immunoprecipitation was performed with appropriate antibodies for 2 h at 4°C. Immunoprecipitated proteins were collected on Protein-G Dynabeads (Novex), separated by SDS-PAGE (NuPAGE, Thermo-Fisher, Waltham, USA) and immunoblots were incubated with appropriate antibodies as indicated and visualised by ECL (GE Healthcare, UK). GST-C/EBPβ constructs and cloning of mutant C/EBPβ proteins were as described previously (Kowenz-Leutz et al 1999; 2010 etc). Antibodies were as follows: anti-C/EBPβ (Leutz lab), anti-C/EBPβ (Santa Cruz; C-19), anti-Flag (Sigma), anti-HA.11 (Covance), anti-TLE3 (Santa Cruz; sc-9124), anti-WDR77/Mep50 (Biomol; A301-562A), anti-Nup50 (Santa Cruz; sc-133859), anti-Mi2 (Santa Cruz; sc-11378; Santa Cruz; sc-11378), anti-MBD2 (Santa Cruz; sc-12444), anti-MBD3 (Bethyl; A302-538A), anti-PRMT5 (Millipore; 07-405), anti-MTA1 (Biomol; A200-280A), DMAP1 (Santa Cruz; B-10), anti-RbbP4 (Abcam), Stat3 (Cell signaling; 9132P), anti-RelA (Santa Cruz; sc-109, anti-GCN5 (H-75; Santa Cruz; SC-20698), anti-ELL (Bethyl; A301-644A), anti-cyclin T1 (Bethyl; A303-499A-M), anti-CDK9 (Bethyl; A303-493A-M), anti-AF9 (Novus; NB100-1565), anti-MLLT1 (Novus; NBP1-26653), anti-PAF1 (Novus; (NB600-274), anti-BRD4 (Bethyl; A301-985A50), anti-AF4 (Santa Cruz; sc-99062), anti-GFP (Roche; 11814460001), anti-SSRP1 (Thermo; PA-22186), anti-SPT16 (Thermo; PA1-12697).

### Homolog mapping and calculation of overlap with known datasets

The two Siersbaek interaction datasets (Siersbaek et al., 2011) are based on the mouse homolog of C/EBPβ and thus all identifiers were mapped to human homologs prior to calculating the overlap between the datasets using the InParanoid Homolog database. PRISMA data was organised into protein groups reflecting the non-unique mapping of peptides to the protein sequences. Within PRISMA data, the unique IPI identifiers served for overlap calculation. Otherwise, the overlap of a set of protein groups (reference dataset) with another dataset was calculated by determining all mutual UniProt identifiers in the two datasets and projecting them onto the protein groups of the reference dataset.

### Calculations of dataset overlaps

Overlap of proteins in the full PRISMA dataset (replicates 1 and 2), the PRISMA core interaction set, and the three other datasets (MudPIT HeLa, IP 3T3L1, and IP SU-DHL1) and their union is shown in the following table. Entries in the diagonal capture the total number of proteins in the dataset.

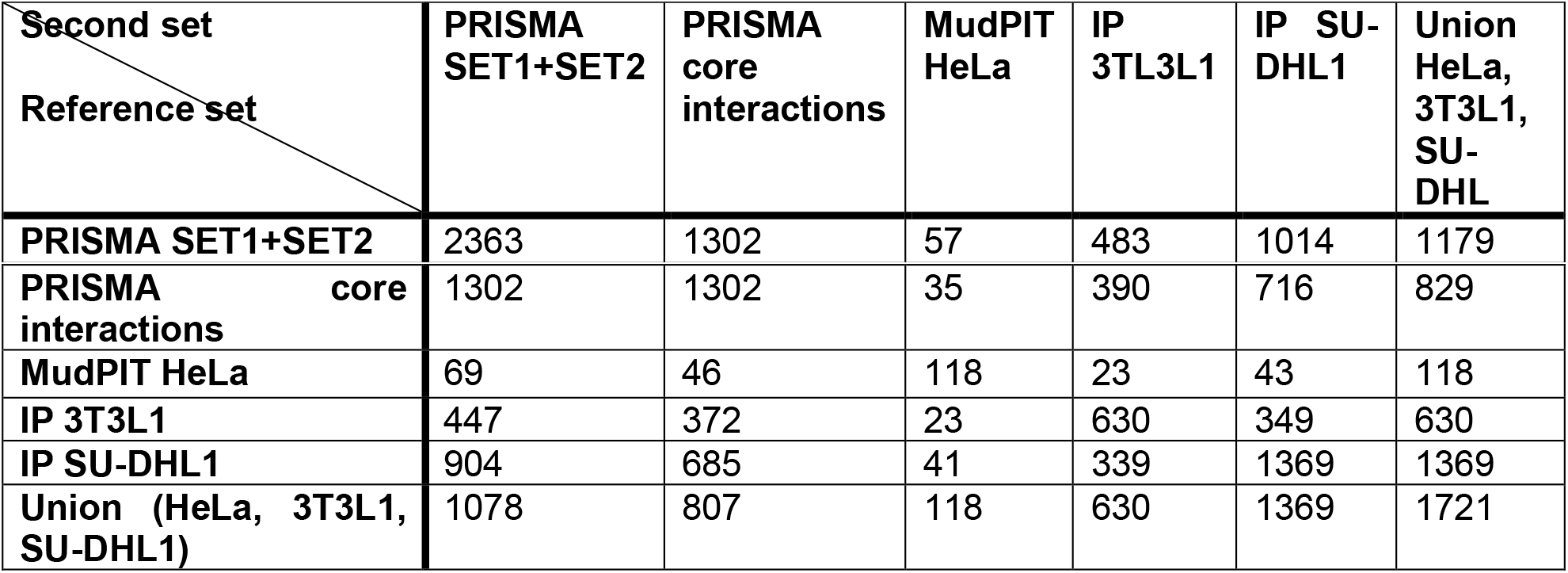

| Second set<br>Reference set | PRISMA<br>SET1+SET2 | PRISMA<br>core<br>interactions | MudPIT<br>HeLa | IP<br>3T3L1 | IP SU-DHL1 | Union<br>HeLa,<br>3T3L1,<br>SU-DHL |
| --- | --- | --- | --- | --- | --- | --- |
| PRISMA SET1+SET2 | 2363 | 1302 | 57 | 483 | 1014 | 1179 |
| **PRISMA core interactions** | 1302 | 1302 | 35 | 390 | 716 | 829 |
| **MudPIT HeLa** | 69 | 46 | 118 | 23 | 43 | 118 |
| **IP 3T3L1** | 447 | 372 | 23 | 630 | 349 | 630 |
| **IP SU-DHL1** | 904 | 685 | 41 | 339 | 1369 | 1369 |
| **Union (HeLa, 3T3L1, SU-DHL1)** | 1078 | 807 | 118 | 630 | 1369 | 1721 |

### FDR calculation

To estimate the false discovery rate (FDR) of protein-protein interaction (PPI) data sets, we employed the method proposed by D’haeseleer and Church (2004) which relies on comparing the intersections of two measured datasets with a reference set to approximate the number of false-positive PPIs in the measured datasets (see **figure below**). We thereby assumed that the reference set and the intersection between the two measured datasets is error-free (i.e. they contain no false-positives).

**Figure:**
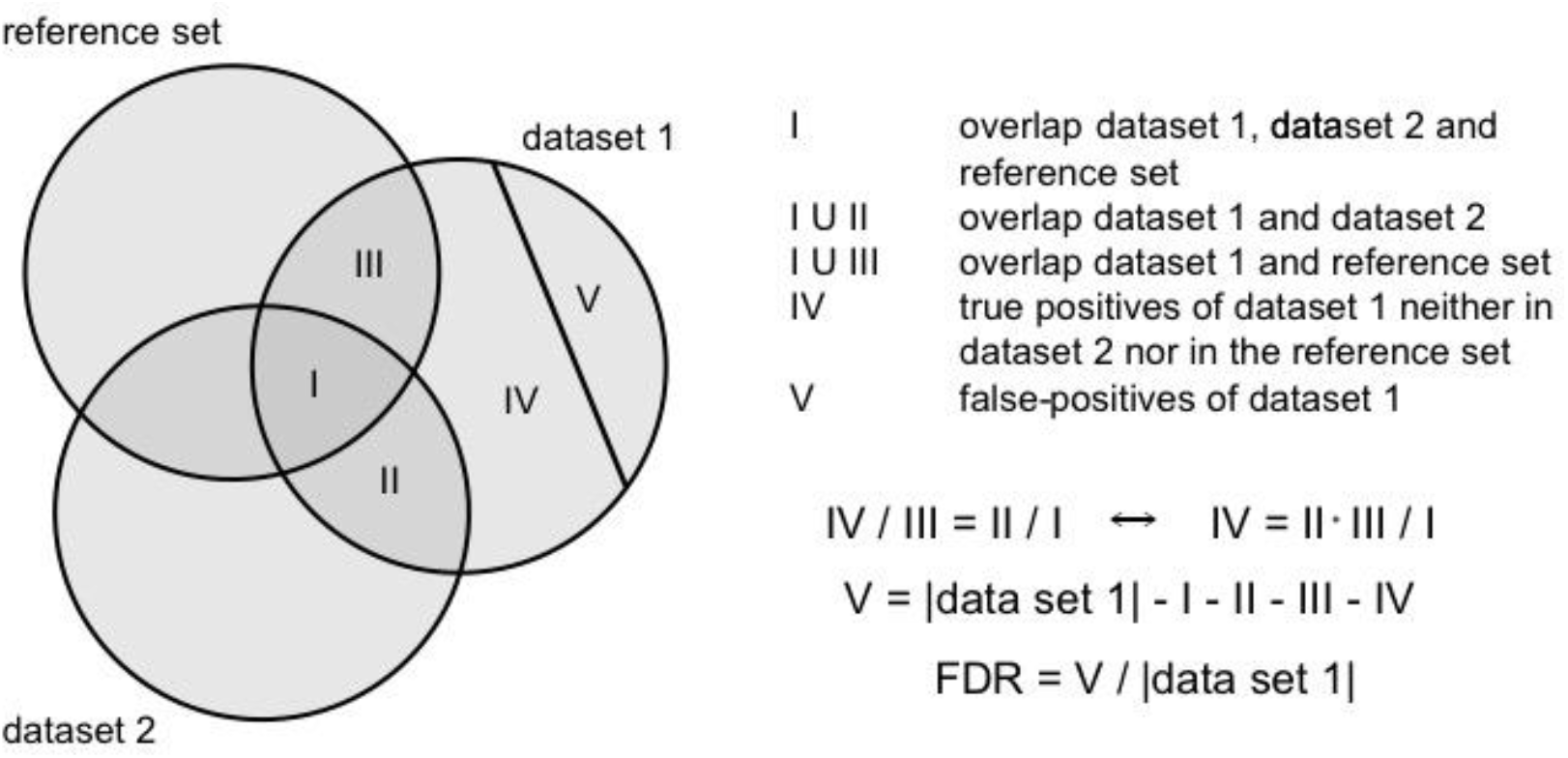
Scheme for the calculation of the false discovery rate (FDR) of dataset 1 according to D’haeseleer & Church (2004). It is assumed that the intersections of the reference set with the datasets (I, III) as well as the intersection between the two datasets (II) are nearly error-free. In addition, the method makes use of the assumption that the reference set overlaps similarly with dataset 1 and dataset 2 (i.e. the relation IV / III = II / I holds. |dataset 1| denotes the number of proteins in dataset 1.

The assumption that the reference set is error-free can be relaxed since we only need to be sure that it contains no bias in how it intersects with the first and second measured datasets and their intersection [D’haeseleer and Church, 2004]. This can be expected if both measured datasets are obtained using the same measurement method, which is the case for PRISMA SET1 and SET2.

Using the calculation procedure described in the **figure above** for PRISMA SET1 and SET2 with the union of the IP SU-DHL1, IP 3TL3L1 and MudPIT HeLa datasets as reference set, we estimated FDRs of 11.2% and 13.9% for SET1 and SET2, respectively. If using as the two measured datasets the restrictions of SET1 and SET2 to proteins also occurring in the PRISMA core interaction set, the FDRs were reduced to 2.5% (SET1 ∩ PRISMA core interactions) and 3.3% (SET2 ∩ PRISMA core interactions). Overlap counts of the PRISMA sets and the reference set (union of IP SU-DHL1, IP 3TL3L1, MudPIT HeLa) which are needed for FDR calculation (i.e. values for the sizes of areas marked by I, II, III, IV, V in the figure above) are given in the following table.

|  | Set size | I | II | III | IV | V | FDR |
| --- | --- | --- | --- | --- | --- | --- | --- |
| **SET1** | 1896 | 862 | 522 | 186 | 113 | 213 | 11.2% |
| **SET2** | 1851 | 862 | 522 | 131 | 79 | 257 | 13.9% |
| **SET1 ∩ PRISMA**<br>**core interactions** | 1203 | 738 | 359 | 51 | 25 | 30 | 2.5% |
| **SET2 ∩ PRISMA**<br>**core interactions** | 1196 | 738 | 359 | 40 | 19 | 40 | 3.3% |

### Ranking of CORUM complexes

We ranked the 1432 complexes obtained from the CORUM database as described in the main text by applying a combination of two criteria: (i) the percentage of proteins of the complex occurring in PRISMA, and (ii) deviation from randomness of the coverage obtained for PRISMA and SU-DHL1. Specifically, for (i), we calculated the percentage for each complex by dividing the number of proteins occurring in the complex from the combined PRISMA replicates 1 and 2 (SET1 + SET2) with matched identifiers using the Perseus tool by the total number of proteins in the complex. For (ii), for each complex, we compared the PRISMA (SU-DHL1) dataset with random protein sets in terms of how many proteins of the complex are covered. Focusing on CORUM data, we considered only the subset of PRISMA and SU-DHL proteins which occurred in the 2678 proteins from the 1432 CORUM complexes, comprising 816 (490) proteins in the PRISMA (SU-DHL) dataset, thus also giving the size of the corresponding random datasets. We first performed the calculations separately for PRISMA and SU-DHL for each of the 1432 complexes. For calculation of a complex of *Cn* >1 proteins, of which *Cp* proteins with 0 ≤ *Cp* ≤ *Cn* occur in PRISMA, we estimated the probability of observing at least *Cp* of *Cn* specific proteins within a random set of size *Pn* = 816 drawn from the total set of *Tn* = 2678 proteins occurring in any of the CORUM complexes. This is a ‘drawing without replacement’ scenario (corresponding to the Fisher’s exact test), for which success is represented by drawing one of a specific subset (of size *Cn*) of proteins, and thus the probability that at least X successes are obtained can be calculated as follows:

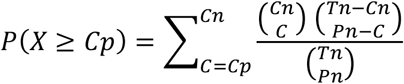

where *Cn* is the maximal number of successes (full coverage of a complex of size *Cn*), *C* is the number of successes (a minimum of *C* = *Cp* proteins of the complex covered because this is how PRISMA or SU-DHL performed, and a maximum of *C*=*Cn*), *Tn* = 2678 (the number of possible results for drawing, which is the number of different proteins in the 1432 CORUM complexes), and *Pn* (the number of draws or size of the random datasets) = 816 (for PRISMA full) and 490 (for SU-DHL1).

For calculation of the probabilities for different complexes, the values of *Cn* and *Cp*, but not of *Tn* and *Pn*, can change. The probabilities correspond to a hypergeometric distribution, and we employed the appropriate *R* base function (dhyper) to compute the values.

The obtained probabilities for each complex, one for PRISMA and one for SU-DHL1, represent how probable it is to obtain the observed (or a more extreme) coverage by chance. We treated the events for PRISMA and SU-DHL1 as independent and multiplied the two probabilities for each complex to obtain the probability that both coverages (or a more extreme) occurred by chance.

We ranked the 1432 complexes based on criterion (i) and (ii) separately. For (i), a high rank corresponded to a high percentage; for (ii), a high rank corresponded to a high significance (i.e. to a low probability; function rank in R, average values for ties). We computed the final ranking from the sum of the two previous rankings (applying minimal values for ties). The complex list is shown together with the probabilities and rankings in Supplemental Table 5.

## Author contributions

Conceptualisation, A.L. and G.D.; Methodology, G.D. and A.L.; Investigation, D.P.H. and G.K.; Validation, E.K.L., R.W., M.Knobl. and M.Kirch.; Resources, G.D., A.L. and U.R.; Data curation, G.D., D.P.H., K.B., M.Kirch., J.W. and A.M.; Writing original draft, A.L.; Writing review and editing, A.L., G.D., E.K.L. M.Kirch., K.B., Visualisation, A.L., G.D., A.M., K.B., E.K.L. and M.Kirch.; Supervision, project administration, and funding acquisition, A.L. and G.D.

## Acknowledgements

We would like to thank Manuel Baesler for generating a web tool (Peptide2Protein) for visualization of PRISMA interactions, Andreas Ladurner, LMU Munich, for FACT expression plasmid constructs, Bernd Lüscher and Juliane Lüscher-Firzlaff, RWTH, Aachen, for MLL complex expression plasmid constructs. Part of the work was supported by DFG Grants to AL, LE770/4-2 and CRC167B11.

---

| enrichment score | pathway / domain description | counts in PRISMA | FDR |
| --- | --- | --- | --- |
| **GO: Biological Processes** |  |  |  |
| GO:0010467 | gene expression | 611 | 4.19e-103 |
| GO:0006355 | regulation of transcription, DNA-templated | 301 | 1.27e-07 |
| GO:0006396 | RNA processing | 267 | 5.63e-128 |
| GO:0007049 | cell cycle | 211 | 3.06e-31 |
| GO:0008380 | RNA splicing | 172 | 7.14e-106 |
| GO:0051276 | chromosome organization | 169 | 1.16e-34 |
| GO:0006461 | protein complex assembly | 128 | 4.17e-12 |
| GO:0016568 | chromatin modification | 89 | 2.50e-13 |
| GO:0051169 | nuclear transport | 86 | 9.46e-38 |
| GO:0006281 | DNA repair | 84 | 9.48e-17 |
| **GO: Molecular Function** |  |  |  |
| GO:0003676 | nucleic acid binding | 745 | 3.8e-199 |
| GO:0003723 | RNA binding | 607 | 0E+00 |
| GO:0005515 | protein binding | 534 | 1.92e-40 |
| GO:0003677 | DNA binding | 269 | 6.43e-17 |
| GO:0005524 | ATP binding | 216 | 2.82e-24 |
| GO:0003682 | chromatin binding | 94 | 4.43e-18 |
| GO:0003712 | transcription cofactor activity | 81 | 2.27e-09 |
| GO:0008134 | transcription factor binding | 66 | 5.86e-09 |
| GO:0019900 | kinase binding | 61 | 3.30e-06 |
| GO:0004693 | cyclin-dependent protein serine/threonine kinase activity | 17 | 6.20e-10 |
| **GO: Cellular Component** |  |  |  |
| GO:0000785 | chromatin | 103 | 6.04e-31 |
| GO:0005681 | spliceosomal complex | 85 | 5.10e-56 |
| GO:0005643 | nuclear pore | 32 | 9.77e-19 |
| GO:0030880 | RNA polymerase complex | 30 | 2.98e-10 |
| GO:0070603 | SWI/SNF superfamily-type complex | 28 | 9.63e-14 |
| GO:0034708 | methyltransferase complex | 24 | 4.86e-09 |
| GO:0016592 | mediator complex | 20 | 3.98e-13 |
| GO:0000178 | exosome (RNase complex) | 12 | 4.26e-09 |
| GO:0016581 | NuRD complex | 11 | 1.50e-09 |
| GO:0016593 | Cdc73/Paf1 complex | 6 | 1.01e-05 |
| **Interpro Domains** |  |  |  |
| IPR027417 | P-loop containing nucleoside triphosphate hydrolase | 102 | 9.71e-30 |
| IPR000504 | RNA recognition motif domain | 80 | 8.77e-53 |
| IPR005225 | Small GTP-binding protein domain | 41 | 2.11e-13 |
| IPR016024 | Armadillo-type fold | 34 | 1.69e-11 |
| IPR017986 | WD40-repeat-containing domain | 32 | 6.78e-10 |
| IPR009072 | Histone-fold | 27 | 3.89e-09 |
| IPR001650 | Helicase, C-terminal | 23 | 1.94e-14 |
| IPR011989 | Armadillo-like helical | 21 | 5.11e-06 |
| IPR011545 | DEAD/DEAH box helicase domain | 20 | 2.11e-13 |
| IPR002041 | Ran GTPase | 20 | 5.34e-12 |
| **PFAM Domains** |  |  |  |
| PF00076 | RNA recognition motif. (a.k.a. RRM, RBD, or RNP domain) | 72 | 4.32e-49 |
| PF00071 | Ras family | 31 | 8.17e-09 |
| PF00400 | WD domain, G-beta repeat | 29 | 1.38e-10 |
| PF00271 | Helicase conserved C-terminal domain | 23 | 2.27e-14 |
| PF00125 | Core histone H2A/H2B/H3/H4 | 23 | 1.89e-09 |
| PF00270 | DEAD/DEAH box helicase | 20 | 2.63e-13 |
| PF00013 | KH domain | 10 | 1.36e-06 |
| PF00244 | 14-3-3 protein | 6 | 2.32e-04 |
| PF01423 | LSM domain | 9 | 2.32e-04 |
| PF00538 | linker histone H1 and H5 family | 5 | 4.50e-04 |

**Supplemental Figure 1.**
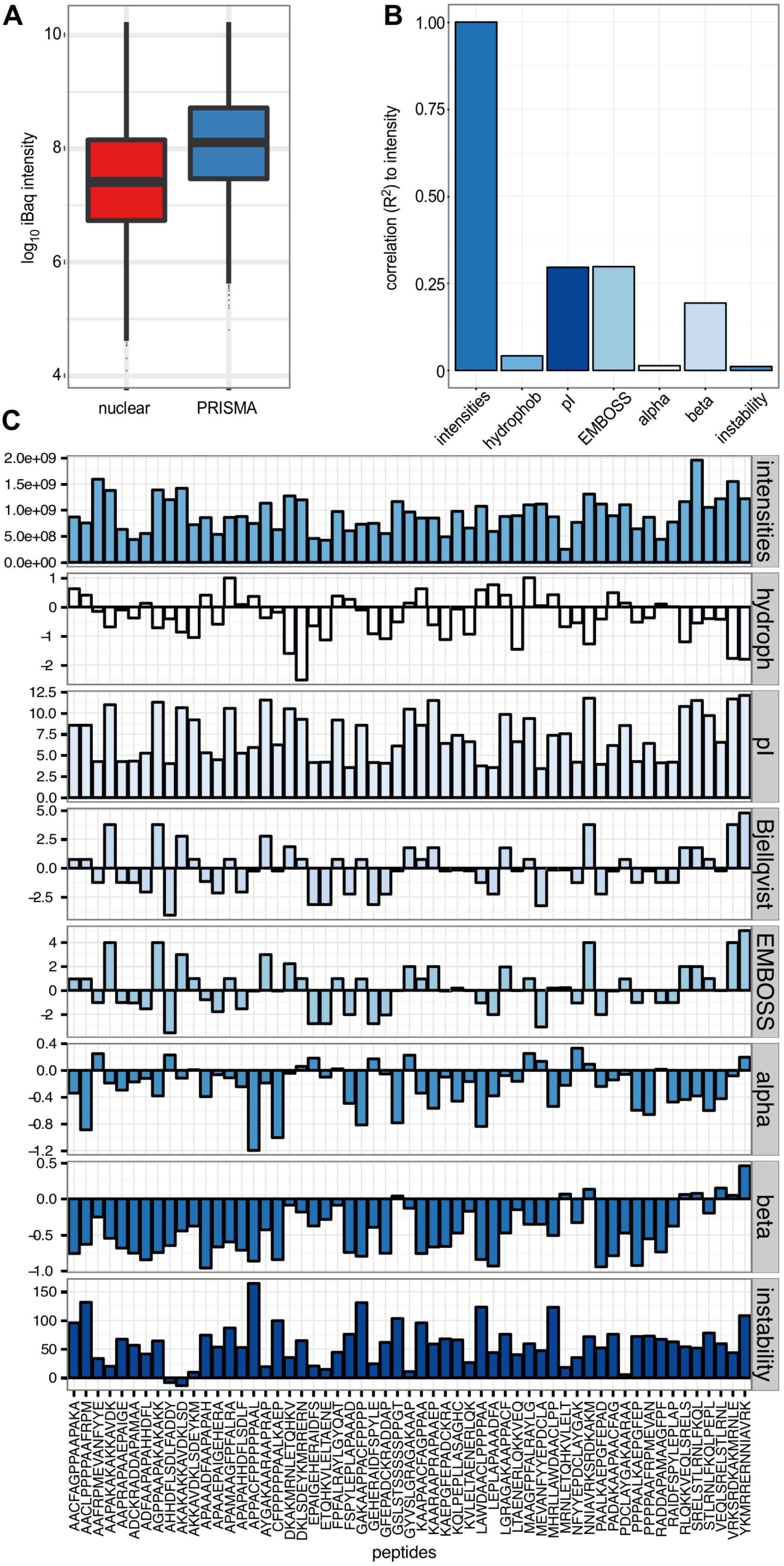
Comparison of enrichment by PRISMA and physico-chemical properties of PRISMA peptides. **A**. Distribution of the copy number estimations, based on the iBAQ quantification for the PRISMA screen and the nuclear extracts. Both, nuclear extracts and PRISMA binders are in the same copy number range. **B**. Correlation of the accumulated intensity of the PRISMA binders to different peptide properties. **C**. Binding of proteins in the PRISMA screen in comparison to various calculated peptide properties. The top panel shows the accumulated binding intensity of the PRISMA binders, while the other panels show the different calculated peptide properties for the peptides listed below.

**Supplemental Figure 2.**
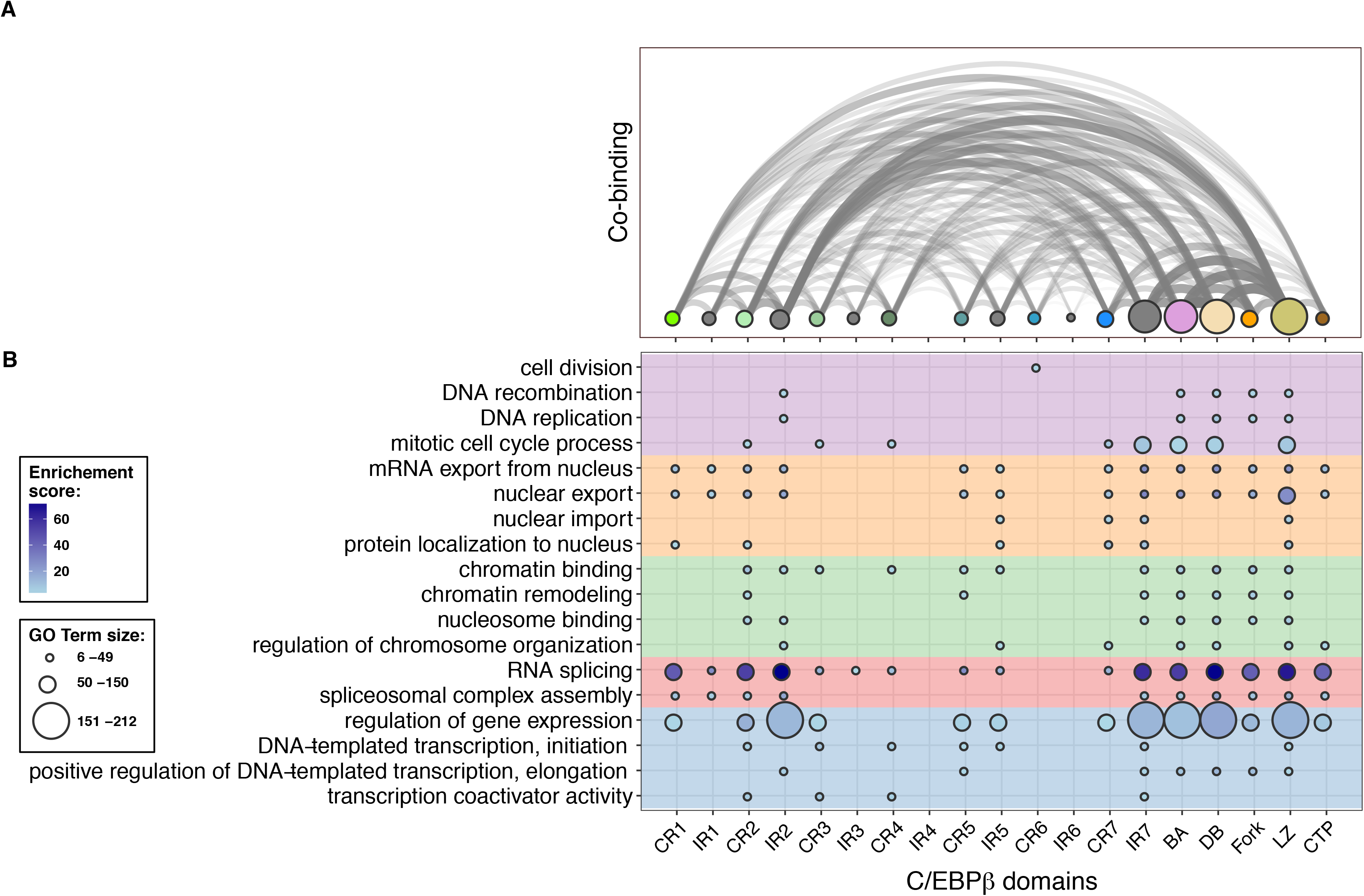
Internal C/EBPβ connections and GO term distribution. **A**. Co-binding of interaction partners to different C/EBPβ regions. The number of interaction partners was calculated that bind to two different regions, as indicated on the abscissa. Dot sizes represent the number of binding partners in the C/EBPβ regions, while the width of the arcs represent the number of interaction partners found in the connected regions. **B**. Enrichment analysis of GO-terms for binding partners in the different regions of C/EBPβ. GO-terms were selected for DNA related processes (pink), nuclear import/export (orange), chromatin (green), RNA splicing (red) and transcription (blue).

**Supplemental Figure 3.**
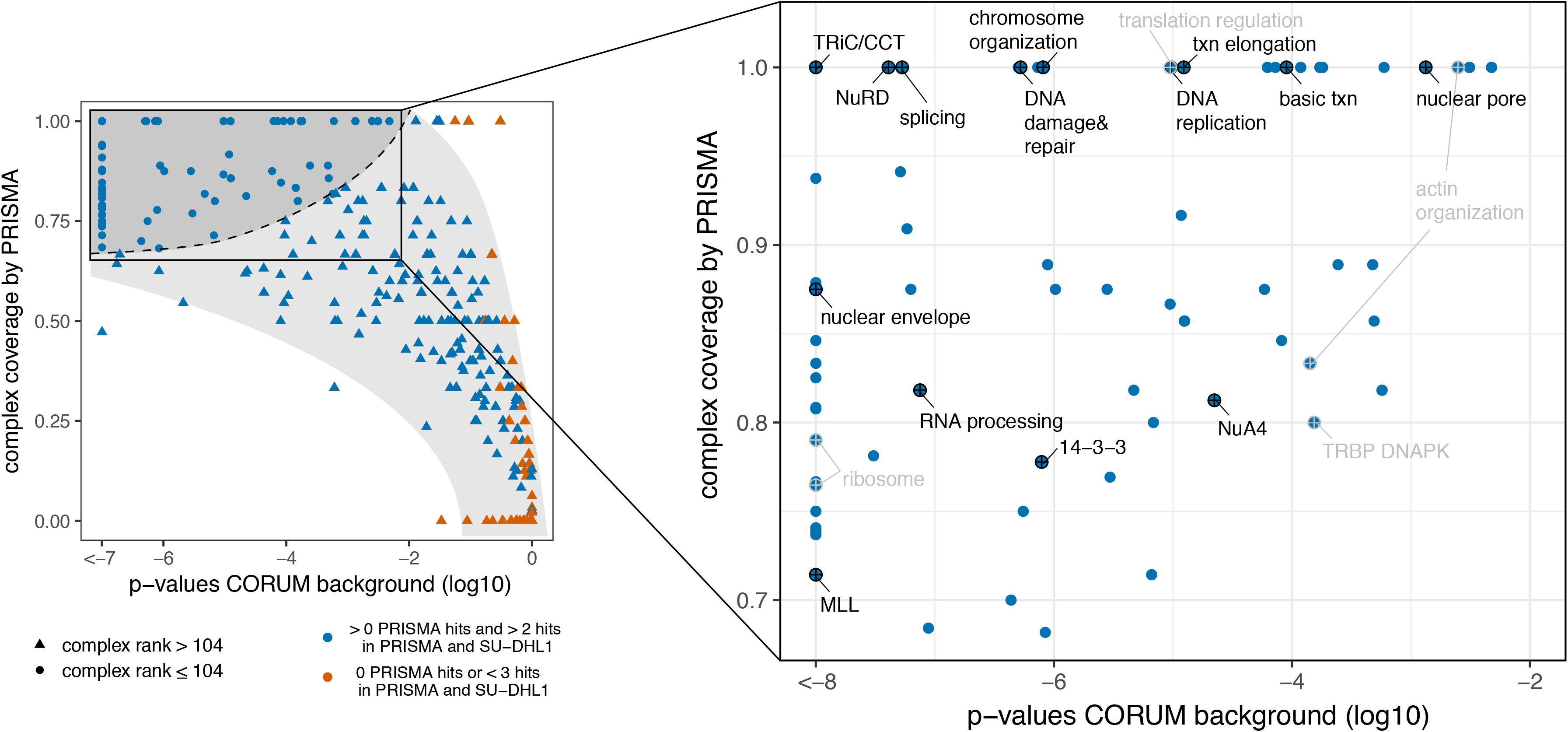
Potential C/EBPβ interacting protein complexes. Complex coverage and p-values for each of the 1432 complexes as used for the ranking of CORUM complexes (**Supplemental Information**). Each dot corresponds to one complex. Red symbols show complexes which have less than three protein hits in PRISMA and SU-DHL1 data sets. Circles represent complexes ranking within the upper quartile (dark grey) (104 complexes, dashed line between upper quartile and lower ranks), triangles encode lower-ranking complexes. The 104 highest-ranking complexes are shown in the close-up on the right. Highest-ranking complexes for each of the 14 categories from Fig. 3B are indicated in black; other complexes which are not included in Fig. 3A in grey.

**Supplemental Figure 4.**
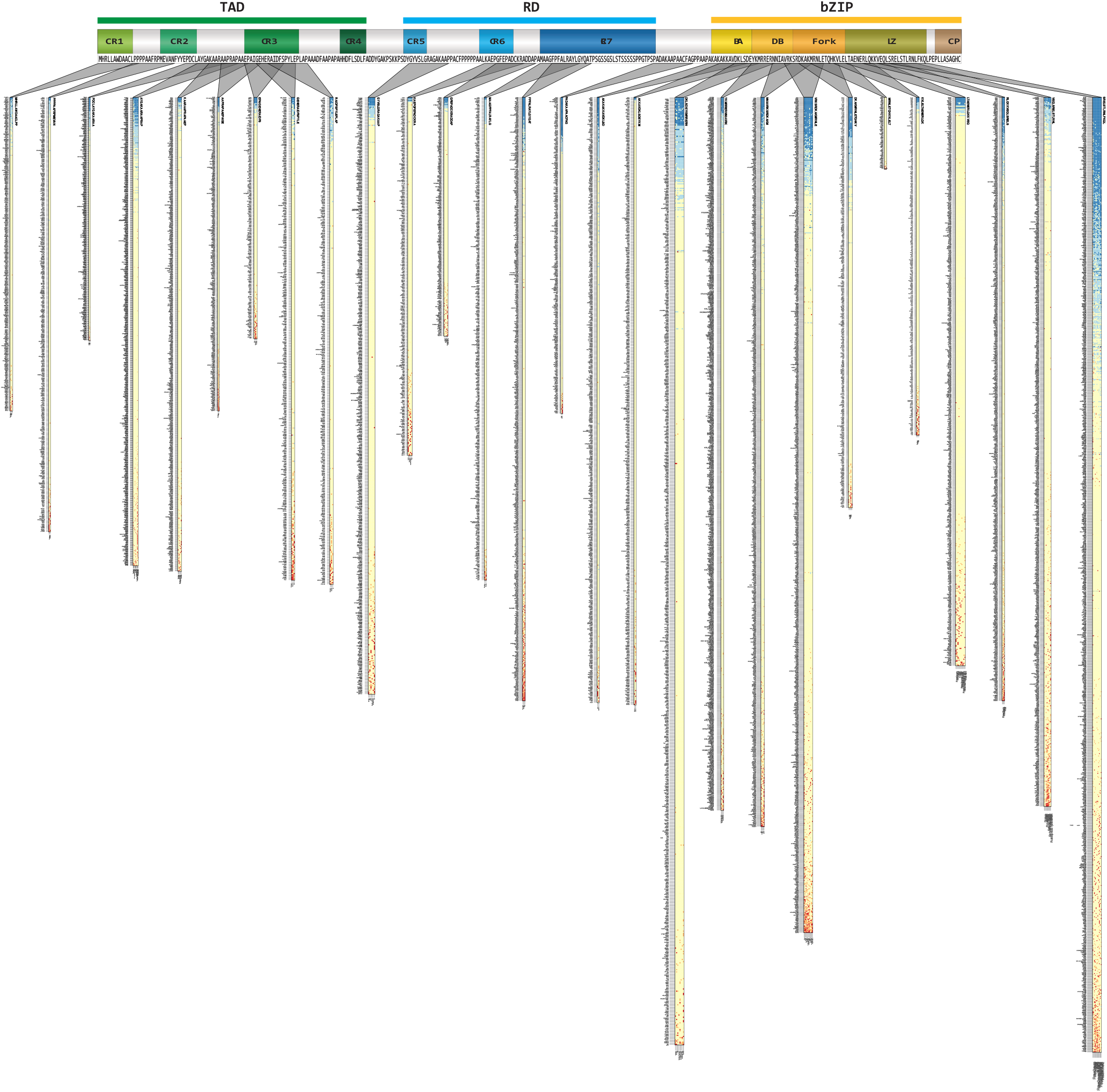
PTM-dependent binding across C/EBPβ. The positions of the PTM-modified peptides are shown on the schematic presentation of the primary C/EBPβ sequence on top. For each peptide, the relative binding to the PTM-modified peptides was calculated in relation to the unmodified peptide. Heatmaps display the relative binding of the proteins on the y-axis in relation to different PTM-carrying peptides, indicated below the heat maps. Red indicates enhanced binding and blue shows reduced binding.

## References

Agarwal, M., Kumar, P., and Mathew, S.J. (2015). The Groucho/Transducin-like enhancer of split protein family in animal development. IUBMB Life 67, 472–481.

Anastasov, N., Bonzheim, I., Rudelius, M., Klier, M., Dau, T., Angermeier, D., Duyster, J., Pittaluga, S., Fend, F., Raffeld, M., et al. (2010). C/EBPbeta expression in ALK-positive anaplastic large cell lymphomas is required for cell proliferation and is induced by the STAT3 signaling pathway. Haematologica 95, 760–767.

Aso, T., Lane, W.S., Conaway, J.W., and Conaway, R.C. (1995). Elongin (SIII): a multisubunit regulator of elongation by RNA polymerase II. Science 269, 1439–1443.

Bhaumik, P., Davis, J., Tropea, J.E., Cherry, S., Johnson, P.F., and Miller, M. (2014). Structural insights into interactions of C/EBP transcriptional activators with the Taz2 domain of p300. Acta crystallographica Section D, Biological crystallography 70, 1914–1921.

Conaway, R.C., and Conaway, J.W. (2013). The Mediator complex and transcription elongation. Biochim Biophys Acta 1829, 69–75.

Cox, J., and Mann, M. (2008). MaxQuant enables high peptide identification rates, individualized p.p.b.-range mass accuracies and proteome-wide protein quantification. Nat Biotechnol 26, 1367–1372.

Cox, J., Neuhauser, N., Michalski, A., Scheltema, R.A., Olsen, J.V., and Mann, M. (2011). Andromeda: a peptide search engine integrated into the MaxQuant environment. J Proteome Res 10, 1794–1805.

Csardi G, N.T.T.I., Complex Systems 1695. 2006. http://igraph.org/ (2006). he igraph software package for complex network research,. http://igraphorg/.

D’Haeseleer, P., and Church, G.M. (2004). Estimating and improving protein interaction error rates. Proc IEEE Comput Syst Bioinform Conf, 216–223.

Di Stefano, B., Collombet, S., Jakobsen, J.S., Wierer, M., Sardina, J.L., Lackner, A., Stadhouders, R., Segura-Morales, C., Francesconi, M., Limone, F., et al. (2016). C/EBPalpha creates elite cells for iPSC reprogramming by upregulating Klf4 and increasing the levels of Lsd1 and Brd4. Nat Cell Biol 18, 371–381.

Disfani, F.M., Hsu, W.L., Mizianty, M.J., Oldfield, C.J., Xue, B., Dunker, A.K., Uversky, V.N., and Kurgan, L. (2012). MoRFpred, a computational tool for sequence-based prediction and characterization of short disorder-to-order transitioning binding regions in proteins. Bioinformatics 28, i75–83.

Dunker, A.K., Garner, E., Guilliot, S., Romero, P., Albrecht, K., Hart, J., Obradovic, Z., Kissinger, C., and Villafranca, J.E. (1998). Protein disorder and the evolution of molecular recognition: theory, predictions and observations. Pac Symp Biocomput, 473–484.

Eaton, E.M., and Sealy, L. (2003). Modification of CCAAT/enhancer-binding protein-beta by the small ubiquitin-like modifier (SUMO) family members, SUMO-2 and SUMO-3. J Biol Chem 278, 33416–33421.

Finn, R.D., Attwood, T.K., Babbitt, P.C., Bateman, A., Bork, P., Bridge, A.J., Chang, H.Y., Dosztanyi, Z., El-Gebali, S., Fraser, M., et al. (2017). InterPro in 2017-beyond protein family and domain annotations. Nucleic Acids Res 45, D190–D199.

Finn, R.D., Coggill, P., Eberhardt, R.Y., Eddy, S.R., Mistry, J., Mitchell, A.L., Potter, S.C., Punta, M., Qureshi, M., Sangrador-Vegas, A., et al. (2016). The Pfam protein families database: towards a more sustainable future. Nucleic Acids Res 44, D279–285.

Gene Ontology, C. (2015). Gene Ontology Consortium: going forward. Nucleic Acids Res 43, D1049–1056.

Hammond, C.M., Stromme, C.B., Huang, H., Patel, D.J., and Groth, A. (2017). Histone chaperone networks shaping chromatin function. Nat Rev Mol Cell Biol 18, 141–158.

He, N., Chan, C.K., Sobhian, B., Chou, S., Xue, Y., Liu, M., Alber, T., Benkirane, M., and Zhou, Q. (2011). Human Polymerase-Associated Factor complex (PAFc) connects the Super Elongation Complex (SEC) to RNA polymerase II on chromatin. Proc Natl Acad Sci U S A 108, E636–645.

Hondele, M., and Ladurner, A.G. (2013). Catch me if you can: how the histone chaperone FACT capitalizes on nucleosome breathing. Nucleus 4, 443–449.

Jeronimo, C., and Robert, F. (2017). The Mediator Complex: At the Nexus of RNA Polymerase II Transcription. Trends Cell Biol.

Jundt, F., Raetzel, N., Muller, C., Calkhoven, C.F., Kley, K., Mathas, S., Lietz, A., Leutz, A., and Dorken, B. (2005). A rapamycin derivative (everolimus) controls proliferation through down-regulation of truncated CCAAT enhancer binding protein {beta} and NF-{kappa}B activity in Hodgkin and anaplastic large cell lymphomas. Blood 106, 1801–1807.

Kajimura, S., Seale, P., Kubota, K., Lunsford, E., Frangioni, J.V., Gygi, S.P., and Spiegelman, B.M. (2009). Initiation of myoblast to brown fat switch by a PRDM16-C/EBP-beta transcriptional complex. Nature 460, 1154–1158.

Kanashova, T., Popp, O., Orasche, J., Karg, E., Harndorf, H., Stengel, B., Sklorz, M., Streibel, T., Zimmermann, R., and Dittmar, G. (2015). Differential proteomic analysis of mouse macrophages exposed to adsorbate-loaded heavy fuel oil derived combustion particles using an automated sample-preparation workflow. Anal Bioanal Chem 407, 5965–5976.

Kovacs, K.A., Steinmann, M., Magistretti, P.J., Halfon, O., and Cardinaux, J.R. (2003). CCAAT/enhancer-binding protein family members recruit the coactivator CREB-binding protein and trigger its phosphorylation. J Biol Chem 278, 36959–36965.

Kowenz-Leutz, E., and Leutz, A. (1999). A C/EBP beta isoform recruits the SWI/SNF complex to activate myeloid genes. Mol Cell 4, 735–743.

Kowenz-Leutz, E., Pless, O., Dittmar, G., Knoblich, M., and Leutz, A. (2010). Crosstalk between C/EBPbeta phosphorylation, arginine methylation, and SWI/SNF/Mediator implies an indexing transcription factor code. EMBO J 29, 1105–1115.

Kowenz-Leutz, E., Twamley, G., Ansieau, S., and Leutz, A. (1994). Novel mechanism of C/EBP beta (NF-M) transcriptional control: activation through derepression. Genes Dev 8, 2781–2791.

Kramer, A., and Schneider-Mergener, J. (1998). Synthesis and screening of peptide libraries on continuous cellulose membrane supports. Methods Mol Biol 87, 25–39.

Krivtsov, A.V., and Armstrong, S.A. (2007). MLL translocations, histone modifications and leukaemia stem-cell development. Nat Rev Cancer 7, 823–833.

Le Guezennec, X., Vermeulen, M., Brinkman, A.B., Hoeijmakers, W.A., Cohen, A., Lasonder, E., and Stunnenberg, H.G. (2006). MBD2/NuRD and MBD3/NuRD, two distinct complexes with different biochemical and functional properties. Mol Cell Biol 26, 843–851.

Lee, S., Miller, M., Shuman, J.D., and Johnson, P.F. (2010a). CCAAT/Enhancer-binding protein beta DNA binding is auto-inhibited by multiple elements that also mediate association with p300/CREB-binding protein (CBP). J Biol Chem 285, 21399–21410.

Lee, S., Shuman, J.D., Guszczynski, T., Sakchaisri, K., Sebastian, T., Copeland, T.D., Miller, M., Cohen, M.S., Taunton, J., Smart, R.C., et al. (2010b). RSK-mediated phosphorylation in the C/EBP{beta} leucine zipper regulates DNA binding, dimerization, and growth arrest activity. Mol Cell Biol 30, 2621–2635.

Leutz, A., Pless, O., Lappe, M., Dittmar, G., and Kowenz-Leutz, E. (2011). Crosstalk between phosphorylation and multi-site arginine/lysine methylation in C/EBPs. Transcription 2, 3–8.

Lichtinger, M., Ingram, R., Hannah, R., Müller, D., Clarke, D., Assi, S.A., Lie-A-Ling, M., Noailles, L., Vijayabaskar, M.S., Wu, M., et al. (2012). RUNX1 reshapes the epigenetic landscape at the onset of haematopoiesis. The EMBO journal 31, 4318–4333.

Luo, Z., Lin, C., and Shilatifard, A. (2012). The super elongation complex (SEC) family in transcriptional control. Nat Rev Mol Cell Biol 13, 543–547.

Lynch, V.J., May, G., and Wagner, G.P. (2011). Regulatory evolution through divergence of a phosphoswitch in the transcription factor CEBPB. Nature 480, 383–386.

Meszaros, B., Simon, I., and Dosztanyi, Z. (2009). Prediction of protein binding regions in disordered proteins. PLoS Comput Biol 5, e1000376.

Minde, D.P., Dunker, A.K., and Lilley, K.S. (2017). Time, space, and disorder in the expanding proteome universe. Proteomics 17.

Mo, X., Kowenz-Leutz, E., Xu, H., and Leutz, A. (2004). Ras induces mediator complex exchange on C/EBP beta. Mol Cell 13, 241–250.

Mohan, A., Oldfield, C.J., Radivojac, P., Vacic, V., Cortese, M.S., Dunker, A.K., and Uversky, V.N. (2006). Analysis of molecular recognition features (MoRFs). J Mol Biol 362, 1043–1059.

Mudunuri, U., Che, A., Yi, M., and Stephens, R.M. (2009). bioDBnet: the biological database network. Bioinformatics 25, 555–556.

Muller, C., Kowenz-Leutz, E., Grieser-Ade, S., Graf, T., and Leutz, A. (1995). NF-M (chicken C/EBP beta) induces eosinophilic differentiation and apoptosis in a hematopoietic progenitor cell line. EMBO J 14, 6127–6135.

Muller-McNicoll, M., and Neugebauer, K.M. (2013). How cells get the message: dynamic assembly and function of mRNA-protein complexes. Nat Rev Genet 14, 275–287.

Nerlov, C. (2008). C/EBPs: recipients of extracellular signals through proteome modulation. Curr Opin Cell Biol 20, 180–185.

Ness, S.A., Kowenz-Leutz, E., Casini, T., Graf, T., and Leutz, A. (1993). Myb and NF-M: combinatorial activators of myeloid genes in heterologous cell types. Genes Dev 7, 749–759.

Orphanides, G., LeRoy, G., Chang, C.H., Luse, D.S., and Reinberg, D. (1998). FACT, a factor that facilitates transcript elongation through nucleosomes. Cell 92, 105–116.

Peterlin, B.M., and Price, D.H. (2006). Controlling the elongation phase of transcription with P-TEFb. Mol Cell 23, 297–305.

Pless, O., Kowenz-Leutz, E., Knoblich, M., Lausen, J., Beyermann, M., Walsh, M.J., and Leutz, A. (2008). G9a-mediated lysine methylation alters the function of CCAAT/enhancer-binding protein-beta. J Biol Chem 283, 26357–26363.

Rappsilber, J., Mann, M., and Ishihama, Y. (2007). Protocol for micro-purification, enrichment, pre-fractionation and storage of peptides for proteomics using StageTips. Nature protocols 2, 1896–1906.

Reimand, J., Arak, T., Adler, P., Kolberg, L., Reisberg, S., Peterson, H., and Vilo, J. (2016). g:Profiler-a web server for functional interpretation of gene lists (2016 update). Nucleic Acids Res 44, W83–89.

Rodier, F., and Campisi, J. (2011). Four faces of cellular senescence. The Journal of cell biology 192, 547–556.

Ruepp, A., Waegele, B., Lechner, M., Brauner, B., Dunger-Kaltenbach, I., Fobo, G., Frishman, G., Montrone, C., and Mewes, H.W. (2010). CORUM: the comprehensive resource of mammalian protein complexes--2009. Nucleic Acids Res 38, D497–501.

Ruthenburg, A.J., Allis, C.D., and Wysocka, J. (2007). Methylation of lysine 4 on histone H3: intricacy of writing and reading a single epigenetic mark. Mol Cell 25, 15–30.

Schwanhausser, B., Busse, D., Li, N., Dittmar, G., Schuchhardt, J., Wolf, J., Chen, W., and Selbach, M. (2011). Global quantification of mammalian gene expression control. Nature 473, 337–342.

Schwartz, C., Beck, K., Mink, S., Schmolke, M., Budde, B., Wenning, D., and Klempnauer, K.H. (2003). Recruitment of p300 by C/EBPbeta triggers phosphorylation of p300 and modulates coactivator activity. EMBO J 22, 882–892.

Sebastian, T., Malik, R., Thomas, S., Sage, J., and Johnson, P.F. (2005). C/EBPbeta cooperates with RB:E2F to implement Ras(V12)-induced cellular senescence. EMBO J 24, 3301–3312.

Shilatifard, A. (2012). The COMPASS family of histone H3K4 methylases: mechanisms of regulation in development and disease pathogenesis. Annu Rev Biochem 81, 65–95.

Siersbaek, R., Nielsen, R., John, S., Sung, M.H., Baek, S., Loft, A., Hager, G.L., and Mandrup, S. (2011). Extensive chromatin remodelling and establishment of transcription factor ‘hotspots’ during early adipogenesis. EMBO J 30, 1459–1472.

Siersbaek, R., Rabiee, A., Nielsen, R., Sidoli, S., Traynor, S., Loft, A., La Cour Poulsen, L., Rogowska-Wrzesinska, A., Jensen, O.N., and Mandrup, S. (2014). Transcription factor cooperativity in early adipogenic hotspots and super-enhancers. Cell Rep 7, 1443–1455.

Slany, R.K. (2009). The molecular biology of mixed lineage leukemia. Haematologica 94, 984–993.

Smith, E., Lin, C., and Shilatifard, A. (2011). The super elongation complex (SEC) and MLL in development and disease. Genes Dev 25, 661–672.

Smits, A.H., Jansen, P.W., Poser, I., Hyman, A.A., and Vermeulen, M. (2013). Stoichiometry of chromatin-associated protein complexes revealed by label-free quantitative mass spectrometry-based proteomics. Nucleic Acids Res 41, e28.

Steinberg, X.P., Hepp, M.I., Fernandez Garcia, Y., Suganuma, T., Swanson, S.K., Washburn, M., Workman, J.L., and Gutierrez, J.L. (2012). Human CCAAT/enhancer-binding protein beta interacts with chromatin remodeling complexes of the imitation switch subfamily. Biochemistry 51, 952–962.

Sterneck, E., Tessarollo, L., and Johnson, P.F. (1997). An essential role for C/EBPbeta in female reproduction. Genes Dev 11, 2153–2162.

Stoilova, B., Kowenz-Leutz, E., Scheller, M., and Leutz, A. (2013). Lymphoid to myeloid cell trans-differentiation is determined by C/EBPbeta structure and post-translational modifications. PLoS One 8, e65169.

Szklarczyk, D., Franceschini, A., Wyder, S., Forslund, K., Heller, D., Huerta-Cepas, J., Simonovic, M., Roth, A., Santos, A., Tsafou, K.P., et al. (2015). STRING v10: protein-protein interaction networks, integrated over the tree of life. Nucleic Acids Res 43, D447–452.

Tompa, P., Davey, N.E., Gibson, T.J., and Babu, M.M. (2014). A million peptide motifs for the molecular biologist. Mol Cell 55, 161–169.

Tsukada, J., Yoshida, Y., Kominato, Y., and Auron, P.E. (2011). The CCAAT/enhancer (C/EBP) family of basic-leucine zipper (bZIP) transcription factors is a multifaceted highly-regulated system for gene regulation. Cytokine 54, 6–19.

Tyanova, S., Temu, T., Sinitcyn, P., Carlson, A., Hein, M.Y., Geiger, T., Mann, M., and Cox, J. (2016). The Perseus computational platform for comprehensive analysis of (prote)omics data. Nat Methods 13, 731–740.

van der Lee, R., Buljan, M., Lang, B., Weatheritt, R.J., Daughdrill, G.W., Dunker, A.K., Fuxreiter, M., Gough, J., Gsponer, J., Jones, D.T., et al. (2014). Classification of intrinsically disordered regions and proteins. Chem Rev 114, 6589–6631.

Weake, V.M., and Workman, J.L. (2012). SAGA function in tissue-specific gene expression. Trends Cell Biol 22, 177–184.

Wethmar, K., Smink, J.J., and Leutz, A. (2010). Upstream open reading frames: molecular switches in (patho)physiology. BioEssays: news and reviews in molecular, cellular and developmental biology 32, 885–893.

Wickham, H. (2009). ggplot2. Elegant Graphics for Data Analysis.

Wickramasinghe, V.O., and Laskey, R.A. (2015). Control of mammalian gene expression by selective mRNA export. Nat Rev Mol Cell Biol 16, 431–442.

Williams, S.C., Baer, M., Dillner, A.J., and Johnson, P.F. (1995). CRP2 (C/EBP beta) contains a bipartite regulatory domain that controls transcriptional activation, DNA binding and cell specificity. EMBO J 14, 3170–3183.

Wright, P.E., and Dyson, H.J. (1999). Intrinsically unstructured proteins: re-assessing the protein structure-function paradigm. J Mol Biol 293, 321–331.

Wright, P.E., and Dyson, H.J. (2015). Intrinsically disordered proteins in cellular signalling and regulation. Nat Rev Mol Cell Biol 16, 18–29.

Xie, H., Ye, M., Feng, R., and Graf, T. (2004). Stepwise reprogramming of B cells into macrophages. Cell 117, 663–676.

Zhang, and Yinghua, L. (2011). The Expanding Mi-2/NuRD Complexes: A Schematic Glance. Proteomics Insights, 79.

